# Reversible Downregulation of HLA Class I in Adenoid Cystic Carcinoma

**DOI:** 10.1101/2024.09.03.610529

**Authors:** Annie Li, Bianca L Gonda, Adam von Paternos, Elizabeth M Codd, Dawn R Mitchell, Markus D Herrmann, Thuc Q Dzu, Prinjali Kalyan, Chengzhuo Gao, Edwin Zhang, Julia J Mendel, Julia C Thierauf, Peter M Sadow, Thomas Denize, Diane Yang, Jong C Park, Florian J Fintelmann, Xin Gao, Ross D Merkin, Atul K Bhan, William C Faquin, Lori J Wirth, Daniel L Faden, Stefan T Kaluziak, A John Iafrate

**Affiliations:** Department of Pathology, Massachusetts General Hospital and Harvard Medical School, Boston, USA; Department of Otorhinolaryngology, Head and Neck Surgery, Massachusetts Eye and Ear and Harvard Medical School, Boston, Massachusetts; Departments of Medicine, Hematology-Oncology, Massachusetts General Hospital and Harvard Medical School, Boston, USA; Department of Radiology, Massachusetts General Hospital and Harvard Medical School, Boston, USA

**Keywords:** MYB, NFIB, MHC-I, B2M, Adenoid Cystic Carcinoma

## Abstract

**Purpose:** Adenoid cystic carcinoma (ACC), a rare and lethal cancer, has shown low response rates to systemic therapies, such as cytotoxic chemotherapy and immune-checkpoint inhibitors (ICIs). Despite numerous clinical trials, some employing aggressive ICI combinations, no effective treatments for patients with recurrent or metastatic adenoid cystic carcinoma have emerged, and ACC mortality rates remain stagnant. Therefore, we aimed to characterize the ACC immune landscape to understand the poor response rates to ICIs.

**Experimental Design:** We leveraged automated multiplex immunofluorescence (mIF), RNA in-situ hybridization, and scRNAseq Gene Expression analysis to identify pathways supporting the cold ACC immune environment and molecularly characterize ACC tumors, adjacent normal tissues, and normal tissues from regions where ACCs arise. In vitro, we treated freshly resected ACCs with interferon-γ or a STING agonist.

**Results:** mIF demonstrated that ACC tumors are immunologically ‘cold’, with few tumor-infiltrating T-lymphocytes (TILs) and low PD-L1 expression. The most striking finding, however, was a very low HLA/B2M class I expression in almost all ACCs, which was reversible through treatment with interferon-γ or a STING agonist. mIF and RNAseq analyses of normal tissues revealed a p63+, NFIB+, basal duct cell population with similarly low HLA/B2M class I expression.

**Conclusions:** Low/absent HLA/B2M expression may explain ACC tumors’ immunologically cold status and lack of response to ICIs. Our findings suggest that the normal cell of ACC origin exists in an HLA-low state, and that pharmacologic manipulation with immune activators, such as STING agonists, can restore HLA/B2M in ACCs, creating a path to urgently needed, effective immunotherapies.

## Introduction

Adenoid cystic carcinoma (ACC) is a rare epithelial cancer of the salivary glands which accounts for approximately 1% of all head and neck cancers,^1^ while less frequently arising in the secretory glands of other organs, including the lacrimal glands, tracheobronchial tree, and breast.^2^ ACC most often exhibits a slow, but progressive disease course, due to its ability to spread through perineural invasion^3,4,5^ and hematogenous dissemination^6,7^ to distant organs. Its three distinct histological subtypes – tubular, cribriform, and solid – represent increasing aggressiveness and decreasing overall survival.^3,8,9,10^ Even with appropriate adjuvant therapy following surgery, local recurrence is common, with increased risk seen with the solid histological subtype, perineural invasion, positive surgical margins, or advanced stage at diagnosis.^11^ Studies have found that 28.5% to 60% of patients with ACC develop distant metastases,^8,12,13,14^ most often to the lungs, but also to liver, bones, and brain, with clinical behavior ranging from indolent to highly aggressive.^13,15,16^ The standard of care for treatment of patients with ACC is surgery and adjuvant radiotherapy.^17^ Systemic therapies in the adjuvant setting are currently recommended only in the context of clinical trials or in cases of metastatic disease.^17,18,19^

Chromosomal translocations, found in approximately 60% of ACC patients,^20^ include the translocation t(6;9), leading to the *MYB*-*NFIB* fusion or rarer t(8;9) leading to *MYBL1*-*NFIB* fusions.^20,21,22,23,24,25^ As these translocations are thought to drive ACC pathogenesis, extensive molecular profiling studies have sought to clarify the mechanisms by which they and other oncogenes, such as *NOTCH1* mutations, drive progression.^3,4,21,26,27^ This work, however, has yet to impact clinical decision-making for patients with targeted therapies for ACC. Approaches based on cytotoxic agents and VEGFR inhibitors,^14,28,19^ as well as immune checkpoint inhibitors (ICI),^14,20,29,30^ have also shown low response rates and are associated with significant toxicities, as well as major impact on quality of life.

Such challenges are highlighted in a 2022 clinical review that examined the outcomes of 55 clinical trials targeting molecular targets for locally recurrent or metastatic ACC.^31^ Approaches to the most common mutational targets and targeting therapies had been designed to: (1) interrupt tumor cell proliferation and survival by targeting c-KIT, EGFR, FGFR, or NOTCH1; (2) prevent neoangiogenesis by targeting VEGF; (3) disrupt immune checkpoint evasion by targeting PD-1; and (4) exploit other targets, such as PRMT5 and ATRA, to develop novel, effective strategies against ACC. None of the included 55 studies was able to yield a complete response in any patient, and partial response was only found in a low fraction of patients in few studies. For example a partial response rate of 16% (5 of 32 patients) was observed in a trial of the multi-kinase inhibitor Lenvatinib; this meager response rate has, however, been the most promising observed, and has led to a category 2B recommendation in the NCCN Head and Neck Cancer guidelines.^18,19,31^ These data illustrate that, despite robust, ongoing efforts, access to effective treatment options for patients with unresectable or recurrent tumors has remained out of reach.

Several challenges have limited the study of the biology of ACC. Chief among them is a lack of validated cell lines, rendering many studies reliant on human tumor tissue biopsies and animal models.^3^ Additionally, the rarity of ACC makes large cohort studies unfeasible, leading to unpredictability in diagnosis and treatment.^32^ However, computational tools now assist discoveries relevant to ACC. A recent proteogenomic study of salivary gland ACCs revealed two molecular ACC subtypes potentially useful for clinical prognostication.^26^ This study showed that ACC-I, with a 37% prevalence, is characterized by a strong upregulation of *MYC* target genes, enrichment of *NOTCH*-activating mutations, and more aggressive disease course. By contrast, ACC-II, with a 63% prevalence and upregulation of both *TP63* and the receptor tyrosine kinases (*AXL*, *MET*, *EGFR*), is associated with a more indolent disease course. Not only were *MYC* and *TP63* sufficient, in this study, to distinguish the two groups, but overlapping, actionable protein/pathway alterations were identified for each subtype.^26^

Similarly, a transcriptomics study of salivary gland ACC evaluated paired normal and tumor tissues from 15 patients.^33^ This study identified a new gene fusion, *TVP23C-CDRT4*, upregulation of *TVP23C* and *CDRT4* in solid ACC tumors, and the well-known *MYB-NFIB* and *MYBL1-NFIB* fusions. In addition, infiltration of T-cells, B-cells, and NK cells in ACC tissues was observed, as was significant upregulation of the nuclear receptor transcription regulator *PRAME*, alongside downregulation of antigen-presenting human leukocyte antigen (*HLA*) genes.^33^

Our ability to analyze the immune environments of rare tumors that respond poorly to immune therapies, such as ACC, has also been greatly improved by automated multiplex immunofluorescence (mIF) platforms, which permit multiparametric studies of single tissue sections.^34^ The COMET automated mIF platform (Lunaphore Technologies SA), which supports staining up to 40 markers on the same tissue slide, confers unique advantages for examining the spatial relationships between different protein markers and their context within different cellular compartments (e.g., normal tissue, tumor and tumor-adjacent stromal tissue, and tumor-infiltrating lymphatic structures). This spatial context is critical for investigating mechanisms that may support the absence of T-lymphocyte infiltrates observed in tumors thereby deemed ‘cold’.^35^

Our study leveraged mIF to explore the immune landscape of ACCs in the spatial tissue context. We sought to better understand why ACCs are resistant to immune checkpoint inhibitors by identifying pathways potentially supporting these tumors’ cold immune environment, as well as molecular characteristics and novel prognostic biomarkers able to differentiate populations of patients with ACC and to advance effective, personalized therapies.

## Results

### ACCs are immunologically ‘cold’

To understand the immune landscape of ACC, we performed a detailed analysis of a cohort of ACC tumors from our pathology archive, utilizing the COMET mIF system, with a panel of 12 antibodies (including T cell, B cell, and macrophage markers, as well as pan-cytokeratin to identify tumor cells, B2M, and Ki67) (Figure 1).The cohort included a total of 27 cases from 26 patients, with 22 ACC samples from 21 patients: 15 head and neck ACCs, two lung (ACC19 and ACC20), two breast (ACC17 and ACC18), two metastases from head and neck ACCs (ACC21 and ACC22), and one ACC from an unknown primary site (ACC7) (Table 1).

**Figure 1.**
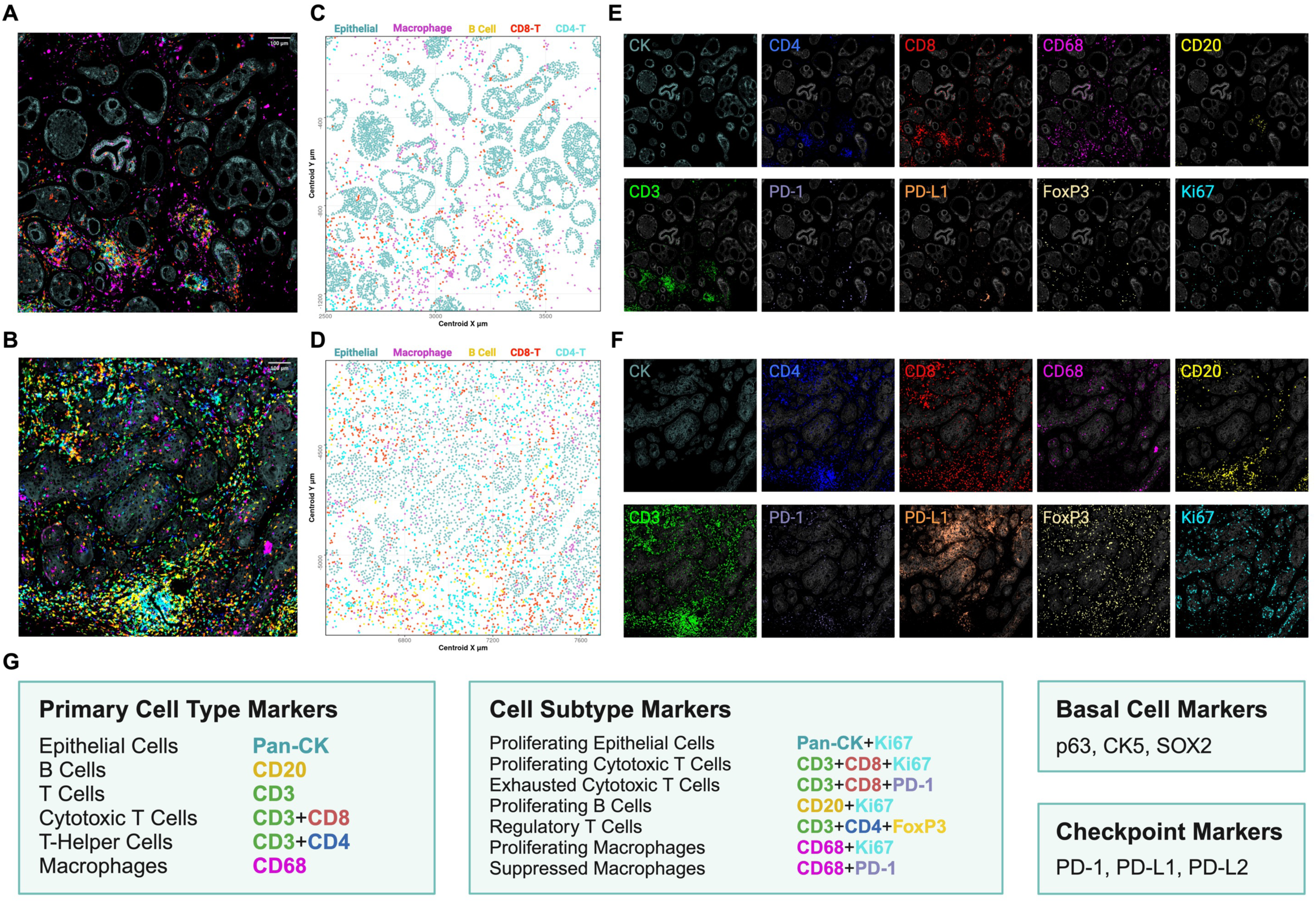
Multiplex Immunofluorescence Panel and Cell Classifications. **A** 10-plex immunofluorescence antibody panel of a representative ACC and **B** SqCC case. **C, D** Maps showing the spatial distribution of classified epithelial cancer cells and immune cells plotted by the x- y- centroid coordinates. Panels in **E** and **F** display expressions of single markers of immune cell markers on the same region of interest. **G** Complete list of protein targets detected by the multiplex immunofluorescence panel. Markers are used for cell classification, cell subtype classification, basal cell detection, and checkpoint marker expression stratification. Adenoid Cystic Carcinoma; SqCC, Squamous Cell Carcinoma; BLC, Basal-like Carcinoma; NB, Normal Breast; NSG, Normal Salivary Gland.

**Table 1.** Demographic, Clinical, and Histological Characteristics of Study Population. Clinical and molecular data for the Adenoid cystic carcinoma patient cohort (n=21) and the control tumors, including Squamous cell carcinomas (n=4) and a basal-like carcinoma of the breast (n=1). *P* values were calculated with a two-sample t-test or a Fisher exact test. *P* value < 0.05 considered significant. H&N, Head and Neck; TNM, Tumor Node Metastasis.

|  |  | Adenoid Cystic<br>Carcinoma Cohort<br>(n=21) | Comparison<br>Cohort<br>(n=5) | P-value |
| --- | --- | --- | --- | --- |
| **Age at Diagnosis** | Median (range) | 50 (35-90) | 67 (48-75) | 0.11 |
| **Gender** |  |  |  | 1 |
|  | Male | 7 | 2 |  |
|  | Female | 13 | 3 |  |
|  | Unknown | 1 | 0 |  |
| **Diagnosis: Adenoid Cystic Carcinoma** |  |  |  |  |
|  | Cribriform | 4 |  |  |
|  | Tubular | 2 |  |  |
|  | Cribriform-tubular | 9 |  |  |
|  | Cribriform/tubular + solid | 2 |  |  |
|  | Solid | 4 |  |  |
| **Diagnosis: Squamous Cell Carcinoma** |  |  |  |  |
|  | Well-differentiated |  | 1 |  |
|  | Moderately differentiated |  | 1 |  |
|  | Non-keratinizing |  | 2 |  |
| **Diagnosis: Basal-like Carcinoma** |  |  |  |  |
|  | High-grade |  | 1 |  |
| **Organ Site** |  |  |  | 0.69 |
|  | Head and neck | 14 | 3 |  |
|  | Lung | 2 | 0 |  |
|  | Breast | 2 | 1 |  |
|  | Metastatic site | 2 | 1 |  |
|  | Unknown | 1 | 0 |  |
| **Stage at Diagnosis** |  |  |  | 0.47 |
|  | I | 9 | 1 |  |
|  | II | 2 | 1 |  |
|  | III | 1 | 1 |  |
|  | IV | 8 | 2 |  |
|  | Unknown | 1 | 0 |  |
| **Status** |  |  |  | 1 |
|  | Alive | 13 | 3 |  |
|  | Deceased | 17 | 2 |  |
|  | Recurrent/Progressive | 11 | 3 |  |
|  | Unknown | 1 | 0 |  |

Additionally, we analyzed a set of five comparator tumors, including four squamous cell carcinomas (SqCC) of the oropharynx and one basal-like carcinoma of the breast. The oral SqCCs were chosen as control tumors, because they, like ACCs, arise in the immune environment of the oral mucosa. The control tumors in this group are generally considered to be ‘hot’. The basal-like carcinoma of the breast serves as a control tumor for ACCs arising from other organ sites, since it is an ACC mimic, showing some similar histological features. Sixteen of 21 ACC patients underwent RNA-based fusion detection, with 12 cases positive for *MYB/MYBL1-NFIB* fusions(Supplemental Table 1). All tumors were reviewed by a Head and Neck subspecialty pathologist (WCF), who verified the diagnoses and ACC histological subtypes. Using a custom image analysis pipeline, we segmented and classified single cells into different cell types based on manually-thresholded values to define individual cell types (Figure 1).

In ACCs, we observed a complex and highly variable tumor immune landscape in the proportion of immune cells (B cells, T cells, and macrophages) to total cells in the tumor area, ranging from 1.5 -34.4%, with a mean of 7.7% (95% CI [4.45%, 10.95%]). This proportion was lower than that of the five non-ACC comparison cohort cases, whose immune cell to total cell ratios were 12.5-22.1%, with a mean of 15.57% (95% CI [10.8%, 20.35%], p-value=0.03) (Figure 2A). When examining the composition of the immune cells, CD8+ cytotoxic T cells showed a significantly lower abundance, at 14.3% (95% CI [10.46%, 18.16%]) in the ACC cases vs. 23.55% (95% CI [10.32%, 36.78%]) in the control tumor cohort (p=0.049) (Figure 2B). A non-significant trend of lower percentages in ACCs versus control tumors could be seen for T-helper cells (24.41%, 95% CI [15.94%, 32.88%] vs. 31.21%, 95% CI [13.87%, 48.55%], p=0.46) and B cells (2.09%, 95% CI [1.09%, 3.1%] vs. 5.5%, 95% CI [-4.93%, 15.93%], p=0.094). CD68+ macrophages represented the only individual cell type showing a trend of greater abundance in ACCs compared to control tumors, comprising 59.18% of immune cells seen in ACC cases (95% CI [48.85%, 69.51%]) vs. 39.73% of immune cells in control tumors (95% CI [5.2%, 74.27%], p=0.12) (Figure 2B).

**Figure 2.**
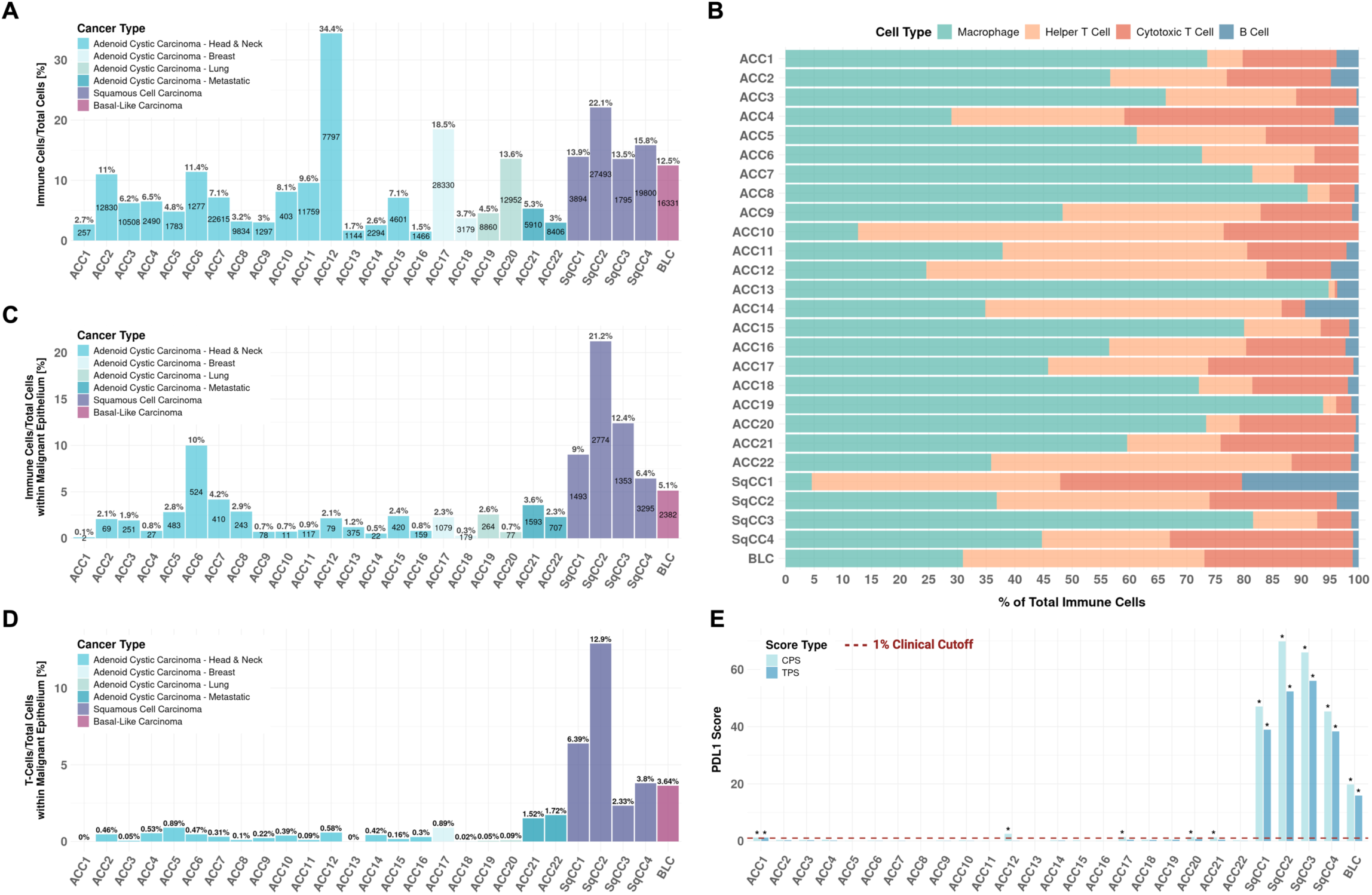
Immune Landscape of ACC Tumors and Comparison Cohort. **A** Percentage of immune cells (macrophages, cytotoxic T cells, T-helper cells, and B cells) to total cells in the tumor area. **B** Relative composition of immune cells. **C** Percentage of immune cells to total cells within the malignant epithelium. **D** Ratio of T cells to total cells within the malignant epithelium. **E** Combined Positive Score (CPS) and Tumor Proportion Scores (TPS) for PD-L1 expression. Cases with scores of TPS >1% and CPS >1 are labeled with *.

These data must be considered in the context that most ACCs arise from the major and minor salivary glands, which are adjacent to the mucosa of the oral cavity and can display an abundance of tissue-resident mucosal immune cells (representative image Supplemental Figure 1A). We were concerned that this proximity had led to overestimation of the tumor-associated immune infiltrate, and thus examined and quantified immune cells within the malignant epithelium itself, limiting our analysis to regions of interest defined by cytokeratin-positive tumor glands (100 glands per case). When examining the amount of total immune cells in the malignant epithelium (Figure 2C), almost all ACC cases showed a lower immune cell infiltration than did the control tumors. Strikingly, and in contrast to the analysis of total T cells in the general tumor sample area, the number of T cells within the malignant epithelium was significantly lower in the ACCs than in the control tumors (Figure 2D). For ACCs, the percent of T cells versus all cells within the malignant epithelium was 0-1.72%, with a mean of 0.42% (95% CI [0.21%, 0.63%]), and for the control tumors was 2.33-12.9%, with a mean of 5.81% (95% CI [0.56%, 11.06%], p<0.001), indicating that ACCs are immunologically cold.

We also observed a difference in the ratio of intra-epithelial B cells between ACCs and tumors (0.017% vs. 0.72%, p=0.012) and intra-epithelial macrophages (1.73% vs. 6.11%, p=0.01). For ACCs, a low percentage of Ki67+ proliferating CD8+ cytotoxic T cells to total T cells was observed in the total tumor area, with a mean of 6.08% (95% CI [3.8%, 8.36%]), while for the control tumors, the percentage was 27.94% (95% CI [0.81%, 55.07%], p<0.001) (Supplemental Figure 1B). These findings indicate a more active, expanding subset of CD8+ T cells in the control tumors than in the ACCs.

### Loss of MHC class I/B2M may underly the lack of ACC immune infiltrate

The lack of tumor-infiltrating T-cells led us to examine several common mechanisms of immune avoidance, specifically: (1) PD-L1/PD-L2 expression on tumor cells, (2) the presence of regulatory T cells in the tumor environment, and (3) MHC class I/B2M expression on tumor cells. The antibody panel on the COMET mIF platform includes PD-L1, whose expression correlates with immune evasion and provides a principal biomarker of immunotherapy response. We used tumor cell fluorescence signals to calculate the Combined Positivity Score (CPS) and Tumor Proportion Score (TPS) for PD-L1.^36,37^ Twenty-one of 22 ACC cases were negative for PD-L1 TPS, with scores of <1% and an average score of 0.22%, CI [0.09%, 0.34%], while the control tumors showed TPS scores ranging from 15.89% to 56% (40.3%, 95% CI [20.73%, 59.83%]) (p<0.0001) (Figure 2E). For the PD-L1 CPS score, 5 ACC cases scored just above the >1 threshold and had an average score of 0.52, while for the control tumors, the CPS scores ranged from 19.76 - 69.8 (49.6, 95% CI [24.83, 74.37]) (p<0.0001) (Figure 2E). PD-L2 is also a ligand of PD-1, but its role in tumor development and progression is still not fully understood. We added PD-L2 to our panel for analysis of a subset of cases, after which 3 out of 10 ACC cases showed a positive CPS score of >1, and 2 cases a positive TPS of >1% (Supplemental Figure 1C). Four of 5 cases in the control tumors showed scores of >1 for CPS and >1% for TPS. These findings suggest that PD-L1 and PD-L2 expression do not have a major role in ACC immune avoidance, consistent with prior publications.^29,38,39^

Since abundant regulatory T cells (Tregs) are known to create an immunosuppressive microenvironment, we quantified the percentage of FoxP3+ regulatory T lymphocytes to total cells. Tregs were sparse in the ACCs (0.45%), whereas this percentage averaged 2.09% for non-ACC cases (p<0.001) (Supplemental Figure 1D). A similar percentage was seen when comparing the number of Tregs to the total number of immune cells in each case: Tregs made up 5.87% of the total immune cells in ACC cases, 95% CI [3.15%, 8.58%]. In contrast, the SqCC cases and the basal-like carcinoma showed an average Treg-to-total immune cells ratio of 12.36%, 95% CI [5.17%, 19.56%] (p=0.041) (Supplemental Figure 1E). When comparing the percentage of regulatory T cells to T-helper cells in ACCs (21.1%) compared to the control tumors (43.39%), we still saw a significantly higher portion in the control tumors (p=0.0035) (Supplemental Figure 1F). Taken together, these findings suggest that Tregs likely do not play a major role in regulating the cancer immune environment of ACCs.

As loss of MHC-I/B2M is a well-described mechanism of tumor immune evasion, we included B2M in our mIF panel. A near complete absence of B2M expression was observed in 20 of 22 ACCs (Figure 3A,B). This contrasted with clear B2M-positivity in four of the five non-ACC control tumors (Figure 3B), with only one SqCC lacking B2M (Supplemental Figure 2A).

**Figure 3.**
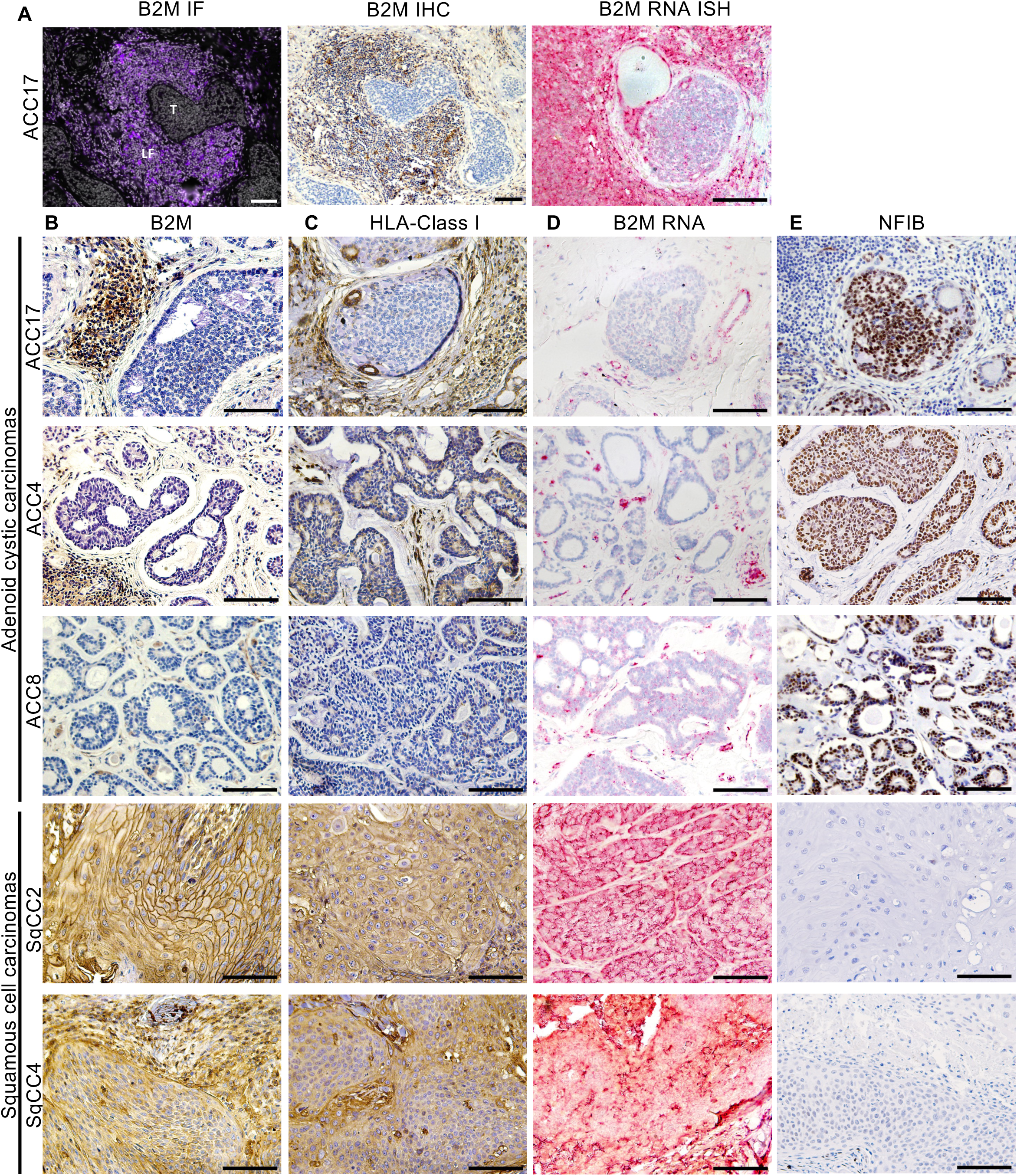
B2M, HLA Class I and NFIB Expression in ACCs and SqCCs. **A** Comparative images for B2M expression obtained by the multiplex immune platform, manual immunohistochemistry, and B2M RNA-ISH on an ACC case showing tumor (T) and a surrounding lymph follicle (LF). **B-E** Histopathology images of ACC and SqCC cases of B2M, HLA-ABC, B2M RNA, and NFIB expression. Representative images are shown at 20x magnification. Scale bar, 100μm. SqCC, Squamous Cell Carcinoma; B2M, beta-2- microglobulin; HLA-ABC, human leukocyte antigen ABC; NFIB, Nuclear factor 1 B-type.

Additionally staining the ACCs with an anti-HLA-A/B/C antibody confirmed that loss of B2M strongly correlated with decreased MHC class I expression (Figure 3C, Supplemental Figure 3A-F). In some ACC cases, these findings showed more positivity and heterogenous expression levels than were seen for B2M, but the levels were still substantially lower than control tumors and adjacent normal tissues (Figure 3C).

Only two ACC cases were identified as having very minimal focal B2M expression within tumor tissue (ACC21 and ACC22). Both cases were metastases, one to a cervical lymph node and the other to the lung, suggesting a unique interaction of the metastatic site microenvironment with tumors cells at the edge of the tumor mass. These two examples of focal B2M expression further suggested that down-regulation might be reversible. The two B2M-positive foci show increased infiltration of T lymphocytes in the tumor glands (Supplemental Figure 2B,C) as well as the highest numbers of T-cells in the malignant epithelium of the ACC cohort (1.52% and 1.72%) (Figure 2D).

### Mechanism of downregulation of MHC class I and B2M in ACCs

B2M can be downregulated in certain tumors through nonsense mutations, increased protein degradation, or decreased transcription.^40,41^ However, examination of published ACC exome sequencing datasets revealed no *B2M* mutations,^42^ suggesting that the lack of B2M expression in the cases we examined were unlikely to have occurred through acquired mutations. As many ACCs can have large amounts of lymphoid tissue and stroma, possibly contributing to *B2M* levels in bulk RNA extracts, we analyzed mRNA expression levels *in situ* at a single cell level using RNA-ISH with a *B2M*-specific probe. In the cancer cells, we observed an absence of *B2M* RNA, whereas a clear maintenance of *B2M* expression levels was seen in the adjacent normal salivary glands and immune cells (Figure 3A,D). This finding strongly suggested that the lack of B2M protein was a result of decreased gene transcription, rather than changes in protein translation or protein stability.

We interrogated the impact of the ACC-associated *MYB-NFIB* fusion on *B2M* transcription. MYB is known to be highly expressed in ACCs because of this fusion, but NFIB expression is not as well studied. We examined 19 of 22 ACC cases in our cohort for NFIB expression levels through IHC, and found NFIB to be highly overexpressed in all 19 ACC cases (Figure 3E).

Since both genes are transcription factors, their high expression suggested one or both might have transcriptional suppressor activity with respect to the *B2M* gene. We searched for potential MYB or NFIB binding sites at the *B2M* promoter, identifying two potential NFIB binding sites between the transcriptional start site and the interferon response element site (Supplemental Figure 4A).

To test whether NFIB was suppressing B2M expression, we knocked down *NFIB* using lentiviral shRNA transduction in five cell lines. Since no confirmed representative ACC cell line models exist, we performed lentiviral shRNA-induced knockdown of *NFIB* in lines with varying levels of NFIB. Three of these cell lines (MCF7, NCI-H446, and S6) showed similar patterns of expression to the ACC cases, with high NFIB and low B2M levels (Supplemental Figure 5A). Knockdown of *NFIB* did not impact B2M protein expression in any line (Supplemental Figure 5B), and overexpression of *NFIB* also did not lead to a change in B2M expression (Supplemental Figure 5C). We also used CRISPR-Cas9 to knock out the two predicted NFIB binding sites at the *B2M* promoter. We performed single cell cloning of cell lines with edits to these sites in three cell lines, MCF7, 293T, and NCI-H446. No increase in B2M protein expression was seen in any of the clones with successful editing at these sites (Supplemental Figure 4B-D). In fact, editing of the predicted site closest to the transcriptional start site resulted in decreased expression of B2M (Supplemental Figure 4B-D).

We also investigated the influence of MYB expression on B2M expression, using siRNA-mediated *MYB* knockdown in MCF7. Western blot analysis revealed no effect of the MYB knockdown on B2M expression (Supplemental Figure 5D). MYB only has two low-confidence binding sites in the *B2M* promoter (JASPAR Score 243 and 249), so we did not proceed with CRISPR/Cas9 editing of that site (Supplemental Figure 5E). Taken together, these data indicate that B2M is not downregulated because of direct transcriptional suppression by either MYB or NFIB.

We explored a second hypothesis, that the ACC cell of origin may have been a B2M-low precursor cell. Reanalysis of a published ACC single-cell RNAseq dataset confirmed that ACC tumor cells are *HLA/B2M* low.^43^ Inspection of the UMAP analysis revealed four clear tumor subclusters, consistent with the known phenotypic variability of ACC tumor cells. One showed myoepithelial/basal markers (*TP63+, NFIB+, ACTA2+*) (cluster C2), while another two showed luminal markers (*KIT*+) (C0, C7) (Figure 4A-D). A fourth subcluster (C3), intermediate between these three clusters, is defined as an *NFIB*-intermediate and *HLA/B2M*-very low state (Figure 4C, D). Reanalysis of normal salivary gland single-cell RNAseq data from the Human Protein Atlas (HPA)^44^ (Figure 4E), and specifically of salivary gland duct cells (Figure 4F), also revealed two *TP63*-high subclusters (C0 and C5), both of which showed significantly lower *B2M* than the *KIT*-high subcluster (p<0.001) (Figure 4G,H). We asked whether we could validate these observations in the glandular ducts in a normal salivary gland sample and a normal breast sample using an expanded 19-plex Lunaphore mIF panel (adding MYB, SOX2, Cytokeratin 5, and p63 to include additional stem cell/basal cell markers), and were able to identify these as p63- and NFIB-high basal cells, that showed very low expression of HLA expression (representative breast duct in Figure 5A-C). In the salivary gland, we identified a corresponding NFIB-high and HLA-low population of basal cells in the salivary gland ducts (Figure 5D-G). Prior reports provide evidence that the basal cells of the intercalated ducts are the cells of origin of ACCs.^45,46^ Detailed analysis of the breast samples revealed a similarly NFIB-high and HLA-low basal cell population in the ducts; in the breast these cells are also smooth muscle actin (SMA)-positive, and of myoepithelial differentiation (Figure 5H-K). Prior reports have also found such a subpopulation of basal cells in the human skin to have a low/negative expression of MHC class I.^47^ These data support a model in which HLA/B2M expression is low in ACCs because the stem cells which transform into ACC naturally show low expression: our data thus indicate that the *MYB-NFIB* fusion might not actively downregulate HLA/B2M expression.

**Figure 4.**
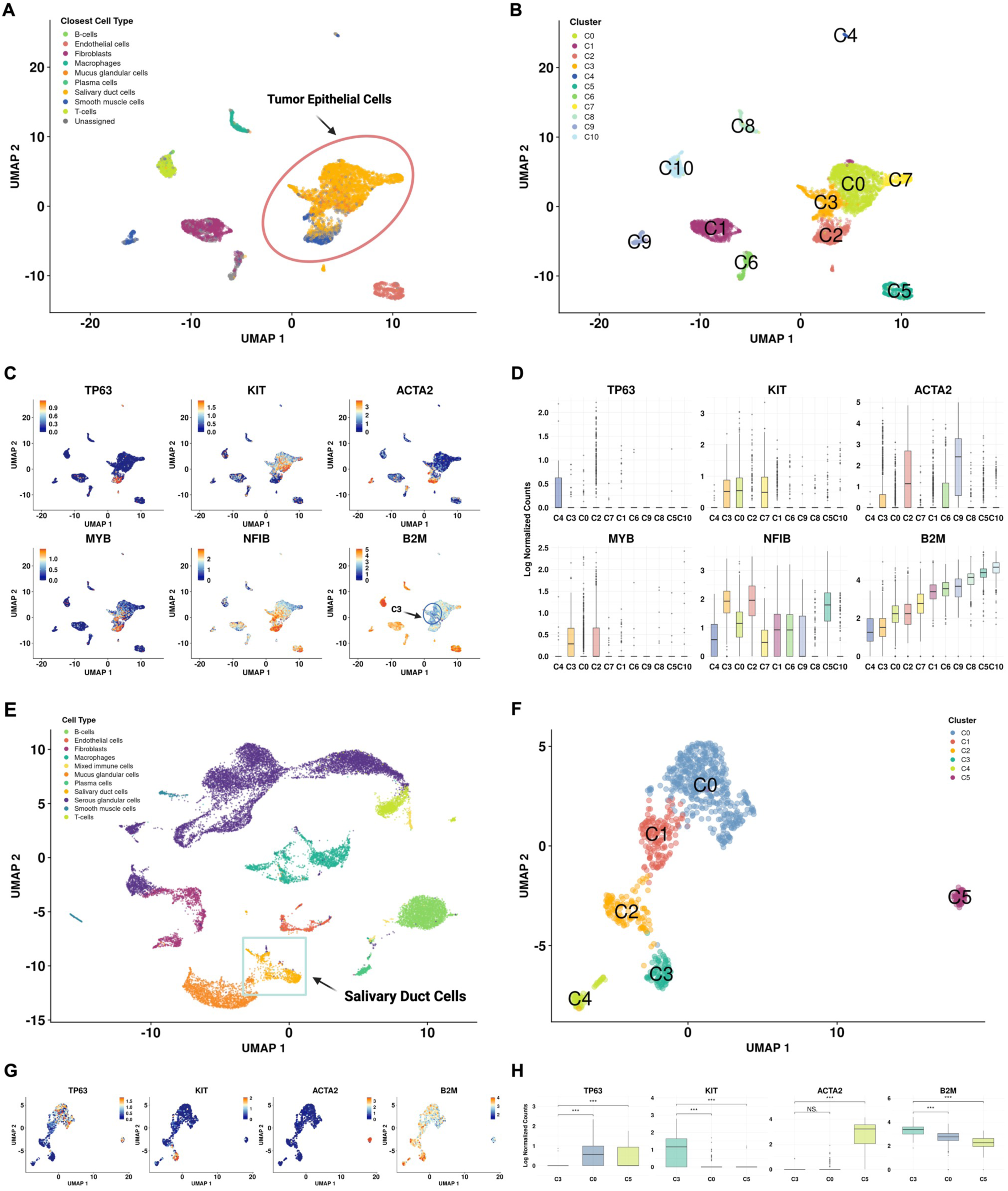
Single cell RNAseq Analysis of Normal Salivary Gland Tissue and ACC Reveal B2M-Low Population of Basal Cells. **A** UMAP representation of reanalyzed scRNAseq data from tumor-only cells of a single ACC case. Cells with matches to known cell types are labeled via color and Tumor Epithelial Cells are circled in red. **B** UMAP of the cells represented in 4A, labeled with cluster IDs produced with Seurat’s Louvain algorithm. **C** UMAPs identifying cells within subclusters (labeled in 4B) with basal-like expression of *TP63+, ACTA2+*, and *NFIB+*, luminal-like expression of *KIT*+, as well as cells with particularly low expression of *B2M* (arrow). **D** Box-plot quantification of transcripts across subclusters, highlighting low *B2M* expression in clusters C3 and C4. **E** UMAP representation of reanalyzed Human Protein Atlas scRNAseq data of normal salivary gland tissue. Cells with matches to known cell types are labeled via color, with Salivary Duct Cells boxed in blue. **F** UMAP of the replotted Salivary Duct Cells highlighted in 4E, labeled with cluster IDs produced with Seurat’s Louvain algorithm. **G** UMAPs identifying cells within subclusters (labeled in 4F), with basal-like expression of *TP63+,* luminal-like expression of *KIT*+, as well as cells with particularly low expression of *ACTA2* and *B2M*. **H** Box-plot quantification of transcripts within salivary duct cells showing *TP63* positive cells have lower expression of *B2M*. ***, *P* < 0.001

**Figure 5.**
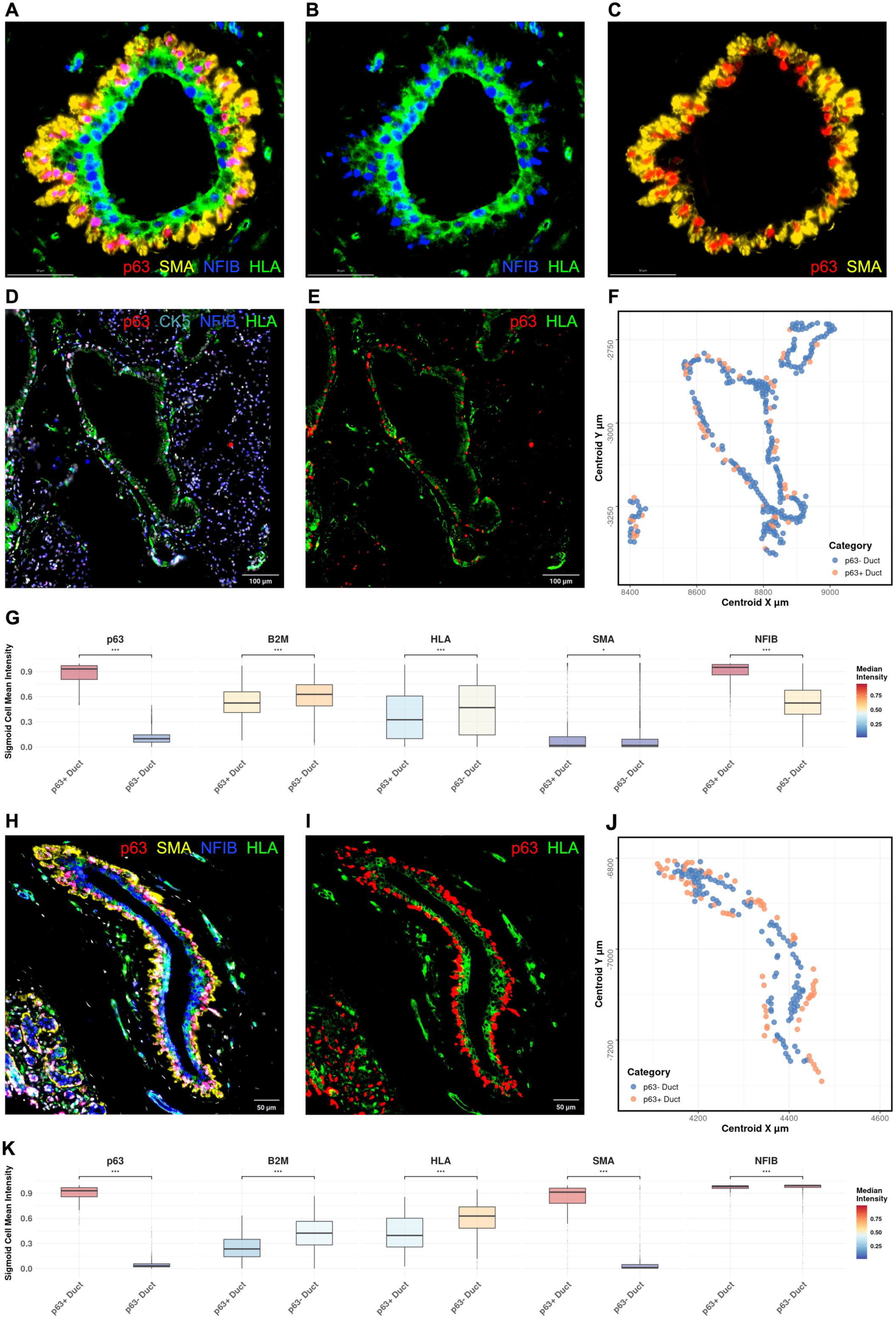
NFIB+/p63+/CK5+ Basal Duct Cells Express Low HLA/B2M. **A** Representative images of a normal breast gland duct with p63, SMA, NFIB and HLA staining, **B** with NFIB and HLA staining and **C** with p63 and SMA staining. **D** Representative image for p63, CK5, NFIB, and HLA-ABC immunofluorescence analysis in a normal salivary gland duct. **E** p63 and HLA-ABC stains highlight low HLA-ABC expression in p63+ basal cells. **F** Spatial distribution of all segmented cells in normal salivary gland ducts differentiating p63+ and p63- cells. Duct depicted in 5A and 5B magnified in top right corner. **G** Box plots showing quantification for all normal ducts on tissue slide. p63+ duct cells are significantly lower in HLA and B2M expression. **H** Representative image for p63, SMA, NFIB, and HLA-ABC immunofluorescence analysis in a normal breast gland duct. SMA and p63 staining highlight myoepithelial/basal cells in pink and yellow. **I** p63 and HLA-ABC stains highlight low HLA-ABC expression in p63+ myoepithelial/basal cells. **J** Spatial distribution of all segmented cells in normal breast gland ducts differentiating p63+ and p63- cells. Duct depicted in 5E and 5F magnified in top right corner. **K** Box plots showing quantification for all normal ducts on tissue slide. p63+ duct cells are significantly lower in B2M and HLA expression. ***, *P* < 0.001.

### Pharmacologic rescue of HLA and B2M in ACC short term cultures

As our mIF data in the two focally B2M-positive metastatic ACC tumors suggested that HLA and B2M low expression could be modulated, we sought to test several pharmacologic approaches known to stimulate the expression of immune genes, including interferon-γ (IFN-γ), STING agonists, and proteasome inhibitors. As no ACC cell line models exist for in vitro drug tests, we employed a short-term culture model to evaluate the effects of in vitro drug treatments on HLA and B2M expression in four freshly resected ACC tissue specimens (Supplemental Figure 6A). These four cases included three *NFIB-MYB* fusion cases and one case with a *NOTCH1* mutation.

Immediately following surgery, the ACC tissue was manually dissected, divided into tissue culture plate wells, and incubated for 48 hours in DMEM/10%FBS with either DMSO, IFN-γ (5ng/ul), the STING agonist ADU-S100 (15mM), pembrolizumab (10 µg/mL), or bortezomib (50 nM). IFN-γ and ADU-S100 both resulted in a strong upregulation of HLA and B2M protein (Figure 6A-C, Supplemental Figure 6B-D). The reintroduction of protein expression was primarily detected at the tumor edges, due to drug diffusion into the tissue. RNA-ISH analysis confirmed extremely strong upregulation of *B2M* transcript levels in the tumor tissue. By contrast, treatments with pembrolizumab or bortezomib led to no significant change in B2M, HLA-ABC expression, or PD-L1 expression (Supplemental Figure 6E-H). IHC analysis of IFN-γ and ADU-S100-treated tumors revealed upregulation of the immune checkpoint protein PD-L1 (Figure 6E). These data from short term cultures indicate that treating ACCs with IFN-γ or STING agonists could restore HLA/B2M expression and possibly result in an anti-tumor immune response. PD-L1 checkpoint induction suggests that any such pharmacologic therapy may need to be combined with anti-PD-1 or anti-PD-L1 antibodies; otherwise, any anti-tumor effect may be self-limiting.

**Figure 6.**
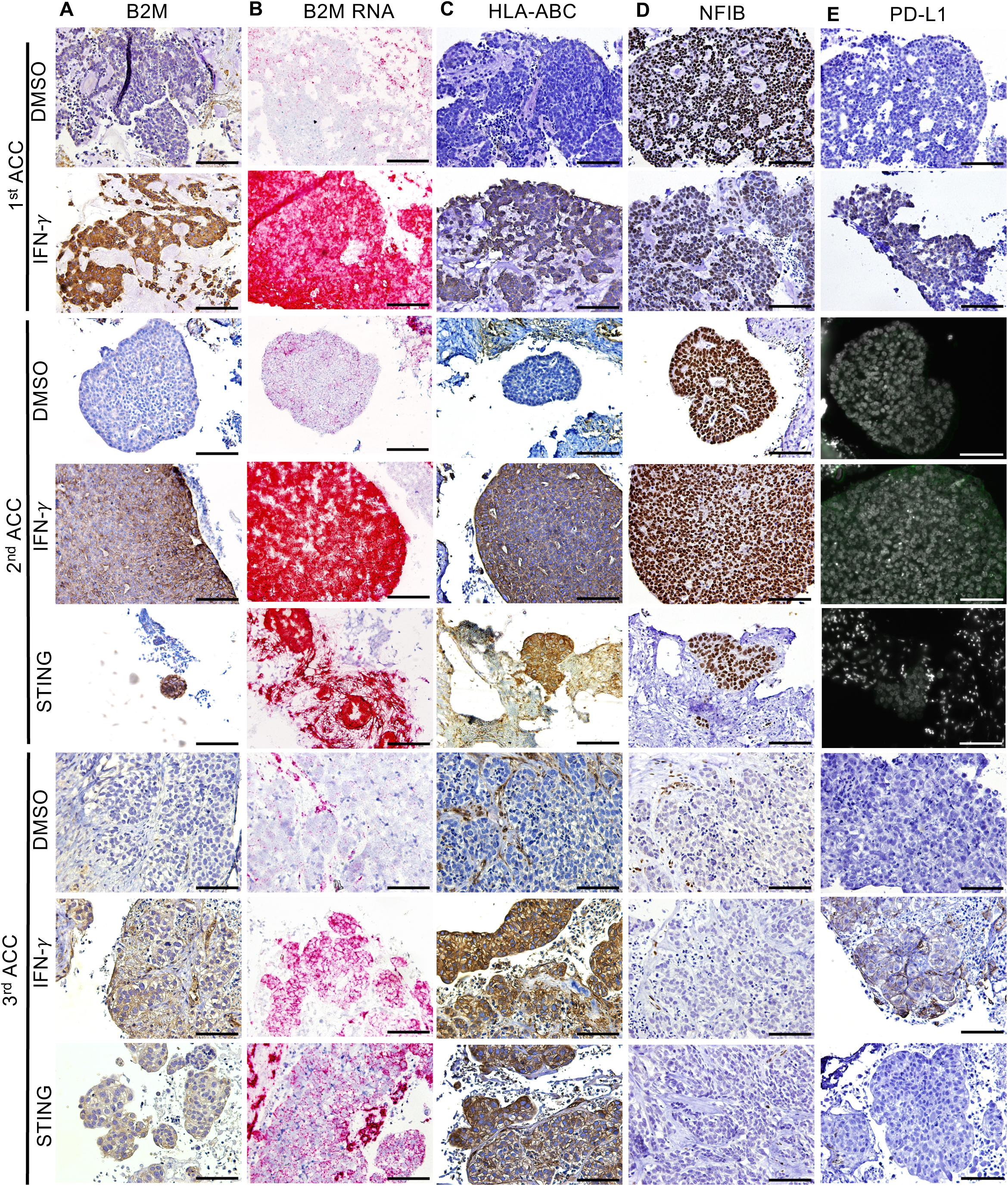
Restoration of B2M Expression with Immune Activator Treatment. **A-E** Histopathology and immunofluorescence images of B2M IHC, B2M RNA levels, HLA-ABC, NFIB, and PDL1 on three ACC slice cultures treated with DMSO (control), interferon- !, or STING agonist. Representative images are shown at 20x magnification. Scale bar, 100μm.

## Discussion

Adenoid cystic carcinoma is typically a slow growing, yet progressive malignancy of the salivary glands and, more rarely, other anatomic sites. In patients with recurrent or metastatic disease following surgery and radiotherapy, treatment options are severely limited.^18^ Despite a number of systemic therapies evaluated to date in clinical trials, activity has been modest at best. The NCCN guidelines recommend clinical trial participation, when available, as a preferred approach for recurrent or metastatic ACC. Other off-label treatment options that may be considered for select patients with recurrent or metastatic ACC include cytotoxic chemotherapy, lenvatinib, or axitinib in combination with avelumab.^19,48^ Therefore, we explored the ACC immune landscape using multiplex immunofluorescence to understand possible reasons that could explain why these tumors are largely unresponsive to immunotherapy. We found that ACCs are immunologically cold and have low expression of PD-L1, confirming several prior studies,^16,20,49,50^ and that this cold phenotype may be due to the downregulation of HLA/B2M, concealing the tumor from T-cell recognition. Using short-term tumor cultures, we further showed that this downregulation is reversible with STING agonists or interferon-γ treatment, potentially revealing new therapeutic approaches to this challenging cancer.

Our study is not the first to describe ACCs as cold tumors. Ferrarotto et al. used RNAseq to evaluate ACCs and found a universal low expression of T-cell checkpoint proteins, a scarcity of tumor antigens, and a clinically-relevant association between the cold ACC tumor microenvironment and higher likelihood of recurrence.^26^ Another 2021 study, using transcriptomic profiles, reported an immune-excluded environment for ACCs, M2-polarized macrophages and myeloid suppressor cells, as well as a low mutational load.^20^ Underscoring the challenges of analyzing rare tumors through scarce biopsy resources and the lack of validated models, another recent study observed a predominantly cold environment with low expression of PD-L1 and an elevation of M2 macrophages.^29^ RNA-sequencing of 62 patient samples found most ACC tumors to be immunologically cold, but about 30% to be ‘hot’; however, spatial analysis revealed a restriction of immune cells to the stroma.^51^ Combined with our data, these studies provide mechanistic insight into why ACCs do not respond to ICI therapy.

The downregulation of HLA/B2M is the predominant immune signal we observed in ACCs; however, we also observed a trend of higher percentages in CD68+ macrophages as compared to control tumors. Since macrophages can have a role in creating an immunosuppressive microenvironment,^52–54^ this is certainly an area worthy of additional investigation. One important observation was that the total number of immune cells in the ACC tumor differed only slightly from the control SqCC tumors; only when measuring T cells within the epithelium itself did we see a striking difference. A likely explanation is that normal salivary glands can have adjacent lymphoid structures, and that the tumor grows into this stromal environment without invoking an anti-tumor response. This explanation is consistent with histologic observations of many tumor glands infiltrating directly into lymphoid follicles. It is unclear why these close approximations do not induce HLA/B2M in the ACC cells.

MHC class I downregulation/B2M loss as a mechanism of immune evasion has been described in a number of viral infections and in cancer.^40,55,56^ Multiple virally-encoded proteins have been shown to directly interfere with expression of HLA/B2M, or to interfere with other components of the antigen presentation machinery.^57,58^ Recent studies have focused on acquired mutations in B2M in tumors resistant to immunotherapies,^57^ and on loss of heterozygosity of the HLA locus in tumors resistant to immunotherapies.^57^ Mutations in HLA/B2M have not been observed in the large number of ACCs studied with exome sequencing.^59^ Our analysis strongly supports a mechanism in which HLA/B2M is downregulated via transcription silencing.

We observed that HLA/B2M downregulation is not linked to the transcriptional activity of the MYB-NFIB gene fusion, as neither knockdown of NFIB and MYB nor CRISPR/Cas9 editing of putative NFIB binding sites in the B2M promoter had any impact on B2M expression. We extrapolated from other NFIB and MYB-expressing tumor lines, since currently no validated ACC cell lines exist. For in vitro drug treatments, the short-term tissue culture provided a sufficient representation of tumor response, similar to that of other models, such as tumor organoids and xenograft models, which are similarly unable to fully recapitulate the human immune response.^60^

Ultimately, ACCs likely do not require active downregulation of HLA/B2M, but instead show a lack of HLA/B2M expression, as do the cells of origin for ACCs, the basal cells of the salivary gland ducts, also known as the duct reserve cells.^45,46^ At baseline, these basal cells exist in an HLA/B2M-low state. Studies have shown that the intercalated duct cells potentially act as transient amplifying progenitor cells and can differentiate into acinar and granular duct cells.^61,62^ scRNAseq and mIF analysis supports these basal cells as stem-like cells, able to give rise to both the myoepithelial and luminal cells that comprise ACCs. These basal duct cells have a neural crest-like expression pattern (NFIB+, SOX2+, p63+),^63–65^ interestingly shared by basal cells of the normal breast ducts. Other cells share this staining pattern, suggesting that other tumors may also have this natural HLA/B2M-low state. We do not know why these NFIB/SOX2/p63 expressing cells have low expression of HLA/B2M, but it likely reflects a general silencing of immune genes in the tumor cells, possibly because of critical signaling pathways, such as NF-KB signaling.

The clinical response of patients with ACCs to immune checkpoint inhibitors has been underwhelming.^66^ In fact, the most recent clinical trials of immunotherapies in this disease have shown very low response rates (2 out of 32 ACC patients), even when using an aggressive combination of ipilimumab and nivolumab.^66^ The study authors showed lower immune cell infiltration in the ACC tumors, the preferential loss of mutational variants with stronger HLA-binding affinity, and a preexisting, clonally skewed intratumoral TCR repertoire, but did not appear to consider the potential role of low MHC Class I in their tumors.

Our data suggest that restoration of HLA/B2M is required in tumor cells before checkpoint inhibition can have a reasonable chance of success. In our study, pharmacologic manipulation of short-term ACC cultures with interferon-γ or a STING agonist each resulted in robust HLA/B2M gene expression and protein upregulation. The B2M promoter has a consensus interferon response element, providing evidence for likely direct transcriptional upregulation of the immune activators via IRF3 or NF-KB. While systemic interferon-γ would be very challenging to implement clinically due to side effects, several STING agonists now in clinical trials target patients with various advanced solid tumors. These studies have largely focused on the ability of STING agonists to modulate the cells of the immune system; our data suggest that perhaps of equal importance is the impact of immune activators on tumor cells. A similar effect of STING agonists on HLA-negative small cell carcinoma of the lung has been recently observed.^67^ Earlier clinical trials involving the STING agonist focused on intra-tumoral injections and modulating the immune response to the tumor. However, the more recent STING agonist trials administer the drug systemically, which seems to be the necessary approach, if the drug needs to modulate the ACC tumor cells themselves, perhaps at metastatic sites. Since STING agonists induce PD-L1 in tumor cells (and likely PD-1 on T cells), checkpoint inhibition will likely remain an important approach in combination with immune activators.

In summary, we report that low/absent HLA/B2M expression explains why ACCs are immunologically cold, potentially explaining their lack of a systemic response to immune checkpoint inhibitors. Our findings suggest that the ability to restore HLA could lead to therapies for treating this particularly difficult to treat tumor, which essentially continues to have no effective systemic treatments despite extensive clinical trials. Future studies should explore the unique concept reported here – that the normal cell of ACC origin exists in an HLA-low state – as well as the possibility that other intractable tumor types may derive from similar HLA-low precursor cells. Finally, we suggest that pharmacologic manipulation of ACCs with immune activators, such as STING agonists, has the potential to restore HLA/B2M in these tumors, and may create a path to urgently needed, effective immunotherapy combinations.

## Materials & Methods

### Patient Cohort and Study Design

This retrospective cohort study was performed with approval from the MGH Institutional Review Board. The study was conducted in accordance with the Declaration of Helsinki. We selected 27 cases from 26 patients who had undergone biopsy or surgery and were diagnosed with ACC, squamous cell carcinoma (SqCC) of the oropharynx, or basal-like carcinoma at the Pathology Department of Massachusetts General Hospital (MGH) between January 2004 and December 2022. Formalin-fixed paraffin-embedded (FFPE) tissue slides, with their associated pathology reports, were obtained from the Pathology archives. The patient cohort included 22 ACC cases (15 head and neck, two metastases from head and neck ACCs, two lung; two breast; one unknown site), 4 SqCCs, and one basal-like carcinoma of the breast.

The SqCC of the oropharynx were chosen to act as control tumors, because these heterogenous tumors are generally considered to be ‘hot’ tumors and arise from a similar immune environment of the oral mucosa as ACCs. The basal-like carcinoma of the breast shows similar histological features as ACCs, and therefore serves as a control tumor for ACCs arising from other organ sites, such as the breast and lung.

### Multiplex immunofluorescence staining and imaging

Multiplex immunofluorescence staining was performed on FFPE tissue slides on the COMET™ platform (Lunaphore Technologies SA). Prior to staining, antigen retrieval and deparaffinization were conducted simultaneously with the PT module (Epredia) by applying the Tris-EDTA based Dewax and HIER Buffer H (TA-999-DHBH, Epredia) at a 1:15 dilution in distilled water for 1 hour at 102 °C. Slides were then washed in Multistaining Buffer (BU07, Lunaphore Technologies) before staining.

Slides were first incubated in Quenching Buffer (BU08, Lunaphore Technologies) for 30 s at 37 °C, next incubated in DAPI for 1 min at 37 °C (1 µg/mL, 62248, Thermo-Fisher Scientific), and then immediately washed in Imaging Buffer (BU09, Lunaphore Technologies). Three rounds of sequential imaging were then performed in DAPI, Cy5, and TRITC channels, with images captured using a 20x objective. After imaging, the slides were washed with a Multistaining Buffer and blocked in 1% BSA (9048-46-8, Research Products International) for 2 min at 37 °C, followed by incubation in two primary antibodies for 8 min at 37 °C. After this 8 min incubation, a 4 min incubation in secondary antibodies Mouse Alexa Fluor Plus 555 (1:200, A32727, Thermo-Fisher Scientific) Rabbit Alexa Fluor Plus 647 (1:400, A32733, Thermo- Fisher Scientific, Waltham, MA), and DAPI was conducted at 37 °C. After a wash in Imaging Buffer, sequential images were again captured. Following imaging, the slides were washed in Multistaining Buffer, and the antibodies eluted by a 2 min incubation in Elution Buffer (Lunaphore Technologies, BU07) at 37 °C. Autofluorescence was then quenched with Quenching Buffer at 37 °C for 30 s. These steps were repeated for the subsequent staining cycles. All primary antibodies are listed in Supplemental Table 2.

### Multiplex Immunofluorescence Image Analysis

#### Image Preprocessing

Leveraging autofluorescence measurements created by the COMET device for the TRITC and Cy5 channels, pixel-based autofluorescence background subtraction was done using HORIZON software (Lunaphore Technologies SA). Subtracting background intensities from foreground staining intensities permits us to more accurately extract the staining intensity for each antibody stain during subsequent analysis steps. These background-subtracted whole-slide images (WSI) were exported as OME-TIFF files for downstream analysis. Additionally, these WSIs were also converted to the Digital Imaging and Communications in Medicine (DICOM) format as VL Whole Slide Microscopy Images. We followed the data model for a DICOM digital pathology workflow,^68^ inserting anonymized clinical metadata as FHIR resources throughout the conversion process and encoding staining procedures, based on TIFF metadata. The DICOM instances were then uploaded to our DICOM store for retrieval according to open DICOMweb API specifications. We leveraged the Slim Viewer,^69^ connected to our DICOM store to visually inspect our datasets, annotate regions of interest for downstream cohort analysis, as well as highlight artifacts to exclude.

#### Cell Segmentation

Cell segmentation was implemented iteratively, leveraging an open-source, pretrained generalist segmentation model, ‘cyto3’,^70,71^ via the cellpose-qupath-extension v0.9.0^72^ in QuPath v0.5.1.^73^ For each WSI dataset, nuclei detection objects were identified via segmentation analysis that leveraged DAPI as the primary channel, with no secondary channel specified. Subsequently, we calculated cell detection objects for each of the cytoplasmic and membrane markers independently by specifying a single marker as the primary channel in conjunction with DAPI as the secondary channel. This process resulted in independent segmentation masks for DAPI, as well as each of the cytoplasmic and membrane markers present in each of the WSI datasets. Next, by intersecting all the centroid coordinates for each independently segmented marker within 130px (∼30µm) of each DAPI segmentation, these data were combined on a cell-by-cell basis to generate complete cell segmentations that included nuclei and cellular contours for all valid cell detections.

#### Cell Fluorescence Measurement Extraction

Intensity summary statistics and cell specific measurements for each valid cell detection were collected post segmentation and cell object reconstruction. Using annotation objects in QuPath, we extracted nuclei and cytoplasmic measurements across optical channels for each antibody staining channel present in each of the WSI datasets. To assess the morphological features of each cell detection, we measured the area, compactness, convexity, and solidity for each nucleus and cytoplasm. To obtain optical measurements within the areas of identified cell detections, for each nucleus and cytoplasm we measured the pixel intensity as a mean, median, 90th percentile, and sum using QuPath and exported these data as a singular csv for each WSI dataset for further analysis in R.

#### Cell Classification

For each sample, a pathologist marked regions of interest, such as total tumor area, malignant epithelium, and normal ducts to be independently analyzed. For each antibody staining channel, the positive expression threshold was manually set based on visual inspection by at least two pathologists. During processing, cells were omitted from analysis according to the following criteria: DAPI below threshold, non-nuclear DAPI (ratio nucleus/cytoplasm < 1), and intensity outliers caused by background subtraction (zeros or extreme values > 10 standard deviations above the mean).

The retained cells were subsequently classified as either positive or negative for each marker by using the manually determined thresholds, and then classified according to the positive marker combination in Supplemental Table 3. These cell classifications and the spatial orientations of their nuclei provided further insight into the imaging of each individual WSI.

#### PD-L1 and PD-L2 Scoring

Cells were classified as PD-L1 or PD-L2 positive or negative based on a digital threshold of the immunofluorescence images, manually set for each sample. The Tumor Proportion Score (TPS) was generated by dividing PD-L1/PD-L2 positive tumor cells by the total number of tumor cells and multiplying by 100. The Combined Positive Score (CPS) was generated by dividing the number of PD-L1/PD-L2 positive cells of any cell type by the total number of tumor cells and multiplying by 100.

### Immunohistochemistry

FFPE slides were deparaffinized and rehydrated via serial incubations in xylene and ethanol. Heat-induced antigen retrieval was performed in a hot water bath in pH 9 Tris/EDTA buffer or pH 6 citrate buffer and stained with primary antibodies anti-beta-2-microglobulin antibody (1:1000, HPA006361 Sigma-Aldrich), anti-beta-2-microglobulin antibody (1:100, MA5-36022, Thermo-Fisher Scientific), anti-HLA-I-ABC antibody (1:1000, ab70328, Abcam), and anti-NFIB (1:100, ab186738, Abcam) for 1 h at room temperature. The corresponding secondary antibody, RTU ImmPress-HRP Goat IgG polymer reagent mouse (MP-7452, Vector Laboratories) or RTU ImmPress-HRP Goat IgG polymer reagent rabbit (MP-7451, Vector Laboratories) was applied for 30 min at room temperature. The sections were then developed using a DAB substrate kit (ab64238, Abcam). The slides were subsequently stained with hematoxylin, dehydrated via serial incubations in ethanol and xylenes, mounted, and cover slipped.

### RNA in-situ hybridization (RNA-ISH)

RNA-ISH was performed using the RNAScope 2.5 assay Reagent Kit- RED (322350, Advanced Cell Diagnostics) according to the manufacturer’s instructions. Sections were deparaffinized and dehydrated with xylene and 100% ethanol before being air dried at room temperature. RNAscope Hydrogen Peroxide was next applied to each section for 10 min and then subjected to manual target retrieval in RNAScope Target retrieval solution for 30 min.

Afterwards, the sections were washed with distilled water, dipped in ethanol, and dried at room temperature. A hydrophobic barrier was drawn around the tissue sections using a barrier pen. Dried slides were covered in Protease Plus treatment and placed into the HybEZ oven (Advanced Cell Diagnostics), set at 40 °C for 30 min. The slides were then washed in distilled water, incubated with an anti-beta-2-microglobulin probe (442201, Advanced Cell Diagnostics) at 40 °C for 2 hours, washed again with the 1X Wash Buffer, and incubated in amplification solutions and Fast-Red detection reagents. Finally, the slides were washed in tap water, stained in hematoxylin and bluing solution, and fully dried at 60 °C before mounting.

### Drug treatment of tissue slice cultures

Surgically resected tumor sections were obtained in cold DMEM (11965, Thermo-Fisher Scientific) containing 10% Fetal Bovine Serum (FBS) (A52568, Thermo-Fisher Scientific) and 1% Penicillin/Streptomycin (15140-122 Thermo-Fisher Scientific), and immediately prepared for in vitro drug treatments. Based on the specimen size, they were manually cut into thin slices with a scalpel and plated into a 12-well cell culture dish. The tissue slices were then incubated in DMEM media, supplemented with 10% FBS and 1% Pen/Strep containing different therapeutic agents in a standard 37 °C humidified incubator with 5% CO2. Tissue slices were subjected either to DMSO (1:1000, D2650, Sigma-Aldrich) as a control^74^ or to one of four different therapeutic agents: 50 nM of bortezomib (S1013 Selleck Chemicals); 10 µM STING agonist ADU-S100 disodium salt (HY-12885A MedChemExpress); 10 µg/mL of pembrolizumab (HY-P9902 MedChemExpress); or 5 ng/mL of interferon-γ (PHC4031, Fisher- Scientific). After 48 h, slices were fixed in 10% neutral buffered formalin for 1 h/mm at room temperature. After fixation they were embedded in paraffin and cut into 5 µm sections for further analysis.

### Cell Culture

All cell lines were obtained from ATCC (Manassas, VA), except for S6, a MGH patient derived glioblastoma multiforme cell line. HEK293T cells, U2OS cells, MCF-7 cells, and S6 cells were cultured in DMEM (11965, Thermo-Fisher Scientific), supplemented with 10% FBS (A5256801, Thermo-Fisher Scientific) and 1% Penicillin/Streptomycin (15140-122, Thermo-Fisher Scientific). NCI-H446 cells were cultured in RPMI-1640 medium (11675, Thermo-Fisher Scientific) supplemented with 10% FBS and 1% Penicillin/Streptomycin. The cells were kept at standard conditions in 37 °C with 5% CO2. The cell lines were tested for mycoplasma infection every two months through PCR testing.

### Total Nucleic Acid Extraction from FFPE Tissue

Regions of interest were circled by the pathologist on an H&E-stained slide from each tissue block, and a corresponding number of 5 µm slides per block needed for extraction were also marked by the pathologist. Total nucleic acid was extracted using the Zymo Quick-DNA/RNA in FFPE MiniPrep Kit (R1009, Zymo Research). FFPE slides were deparaffinized in xylene for 5 min, followed by a wash in 100% ethanol for 5 min. Using a scalpel blade, the region of interest was then manually scraped directly into an Eppendorf tube containing 95 µL of DNase/Rnase free water, 95 µL 2X Digestion Buffer, and 10 µL Proteinase K. Each Eppendorf tube was incubated for 1 hour at 55 °C, and then for an additional 20 min at 94 °C. 600 µL of DNA/RNA Lysis Buffer was next added to the tissue and centrifuged at 10,000 rpm for 60 seconds. Following this, the lysate was washed with 100% ethanol and transferred into a Zymo-Spin IICR Column and collection tube. Nucleic acid was extracted from the lysate according to the manufacturer’s instructions, and DNA/RNA concentrations were quantified using the Qubit RNA HS Assay kit (Q32855, Thermo-Scientific) or Qubit ssDNA HS Assay kit (Q32854, Thermo-Fisher Scientific).

### NFIB and MYB knockdown

NFIB shRNA lentiviral particles (sc-43565-V, Santa Cruz Technologies) or Control shRNA Lentiviral Particles (sc-108080, Santa Cruz Technologies) were used for lentiviral transduction in cell lines on 12-well plates at 60% confluency, according to the manufacturer’s protocol.

After 24 h of transduction, the cells were selected with puromycin (A1113803, Thermo Fisher) for 48h at the following concentrations: MCF7 2ug/ml, S6 2ug/ml, 293T 3ug/ml, U2OS 1.5ug/ml, and NCI-H446 0.75ug/ml. For the c-MYB shRNA knockdown, the cells were seeded in 6-well plates and transfected either with the ON-TARGETplus human MYB siRNA SMARTPool (L-003910-00-0005, Horizon Discovery) or the Control ON-TARGET non-targeting negative Pool (D-001810-10-05, Horizon Discovery) at a concentration of 90nM for 48 h and 72 h per manufacturers protocol using the DharmaFECT 1 Transfection Reagent (T- 2001-01, Horizon Discovery).

### Lentivirus Production

Lentivirus was produced in 293T Lenti-X cells (632180, Takara) with Roche Xtreme-Gene Transfection reagent (XTG9-RO, Millipore Sigma), the packaging plasmid psPAX2, and the VSV-G expressing plasmid pMD2G (#12260, #12259, Addgene). The viral supernatants were collected after 48 h of transfection. The target cells were transduced in media containing 5 ug/ml of polybrene (SC-134220 Santa Cruz technologies) for 24 h and afterwards selected with puromycin for 48 h.

### Plasmid Construction

Two guide RNAs (gRNAs) targeting the NFIB binding site on the B2M promoter region gRNAa: AAACGCGTGCCCAGCCAATC, gRNAb: CTGGCACTGCGTCGCTGGCT were cloned into the plasmid LentiCrispr V2 (plasmid #52961, Addgene) NFIB overexpression plasmid utilized the NFIB full-length sequence in the pLV plasmid backbone.

### Western Blot Analysis

Western blotting was performed by applying standard protocols. The primary antibodies were diluted in 5% milk at the following dilutions: anti-NFIB (1:1000, ab186738, Abcam), anti-c-MYB (1:500, 12319, Cell Signaling), anti-B2M (1:500, MA5-36022, Invitrogen), and anti-Cyclophilin B (1:5000, 43603S, Cell Signaling) and then incubated for 1 h. The secondary antibodies – anti-rabbit IgG, horseradish-peroxidase-linked (1:2000, 7074S, Cell Signaling) anti-mouse IgG, horseradish-peroxidase-linked (1:2000, 7076S, Cell Signaling) – were applied for 1 h. The membranes were then developed with SuperSignal™ West Pico PLUS Chemiluminescent Substrate (34580, Thermo Scientific), and imaged using autoradiographic film (XAR ALF 2025, Lab Scientific).

### scRNAseq Gene Expression Analysis

The normal salivary gland scRNAseq raw count table and cell type annotations were downloaded from Human Protein Atlas (HPA)(proteinatlas.org). Cells with fewer than 500 genes or 1000 reads, greater than 40% ribosomal reads, or greater than 30% mitochondrial reads were removed. scRNAseq read counts for a primary salivary gland ACC tumor were downloaded from NCBI Gene Expression Omnibus accession GSE217084 and filtered in the same manner. The filtered counts were processed and analyzed independently for both datasets using Seurat v5.0.2. Briefly, the counts were log-normalized and scaled. Using the first 20 principal components of the top 2000 most variable genes, the cells were then clustered using Louvain modularity optimization of the shared nearest neighbors (SNN) graph and visualized using Uniform Manifold Approximation and Projection (UMAP) dimensionality reduction.

The subset of cells labeled as salivary duct cells in the HPA normal salivary gland dataset were also re-clustered and visualized separately by using the same parameters. For the ACC primary tumor dataset, cell types were labeled using CHETAH v1.18.0, with a custom reference generated from the HPA data processed as described above. All expression-valued UMAPs were generated using the ‘FeaturePlot’ function from Seurat, with minimum and maximum cutoffs set to the 0.01 and 0.99 quantile respectively for each gene shown. Box plots with significance annotations were generated using ggplot2 v3.5.0 and ggsignif v0.6.4.

### Fusion Detection

To assess MYB-NFIB or MYBL1-NFIB fusion positivity in ACC cases, approximately 200ng of total RNA were extracted from FFPE tissue identified as tumor by a molecular pathologist, and then processed according to vendor specifications with the ArcherDX FusionPlex Pan Solid Tumor v2 kit (AB0137, Archer - Integrated DNA Technologies). Final sequencing libraries of samples were multiplexed and sequenced on the Illumina NextSeq 550 platform with the High Output v2.5 (300 Cycle) kits (20024908, Illumina). Raw sequencing data were demultiplexed and converted to fastq files with BCL Convert v4.2.1, which were subsequently uploaded to the Archer Analysis version 7.1.0-14 (Archer - Integrated DNA Technologies). Additionally, we leveraged a custom fusion analytics pipeline, implemented in Bash, to quality control raw fastq reads with TrimGalore,^75^ align reads to the GRCh38 reference assembly^76,77^ with STAR-Fusion,^78^and identify fusions using Arriba.^78^ Fusion results for each sample were verified on a case-by-case basis by expert molecular pathologist review (AJI).

## Statistical Analysis

Differences of means between continuous variables for ACC vs. the control tumors were compared using Student’s t-test (‘t.test’ in R). Fisher’s exact test (‘fisher.test’ in R) was used to assess differences in proportions between categorical metadata features. For both scRNAseq and COMET data, Mann-Whitney U tests were performed to assess the locational shift of expression distributions between cell subsets of interest via the ‘wilcox.test’ function in R. Prior to statistical analysis and box plot visualization, all COMET cell mean intensities were transformed using the following sigmoid function to obtain values ranging from 0 to 1 with the threshold mapped to 0.5:

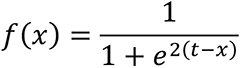

where *x* is the observed cell mean intensity and *t* is the intensity threshold for a given marker. This transformation allowed us to minimize the effects of outliers and to better visualize differences between values in proximity to the threshold. All statistical calculations were performed using base R v4.3.1.

## Data Availability

All code used to perform statistical analysis, as well as relevant figures is available at: https://github.com/IafrateLab/li_et_al-2024.

## Acknowledgements

We thank our colleagues in the clinical teams at Massachusetts Eye and Ear Hospital and Massachusetts General Hospital for the assistance provided with tumor collection. Figure 1, 2, 4, 5, and Supplemental Figures 1 and 6 were partially created with BioRender.com.

## Statement of Translational Relevance

When ACC reaches an advanced stage, palliative support is all oncology can currently offer. ACC thus represents a major unmet clinical need. Two novel findings from our studies of the ACC immune environment have important translational relevance. First, we identified a striking downregulation of HLA Class I and B2M expression in all ACC cases. We also identified a subpopulation of p63+/NFIB+ basal duct cells having similarly low HLA/B2M class I expression patterns, suggesting that ACCs arise from basal duct cells and maintain their low HLA/B2M expression patterns. Second, in vitro treatment with interferon-γ or a STING agonist led to strong upregulation of HLA/B2M class I in fresh ACC slice cultures, showing that HLA/B2M downregulation can be reversed to help restore antigen presentation and adaptive immune system recognition. Additionally, PDL1 expression was upregulated, pointing towards an urgently needed, potential STING-ICI combination therapy for ACC patients with advanced disease.

---

**Supplemental Figure 1.**
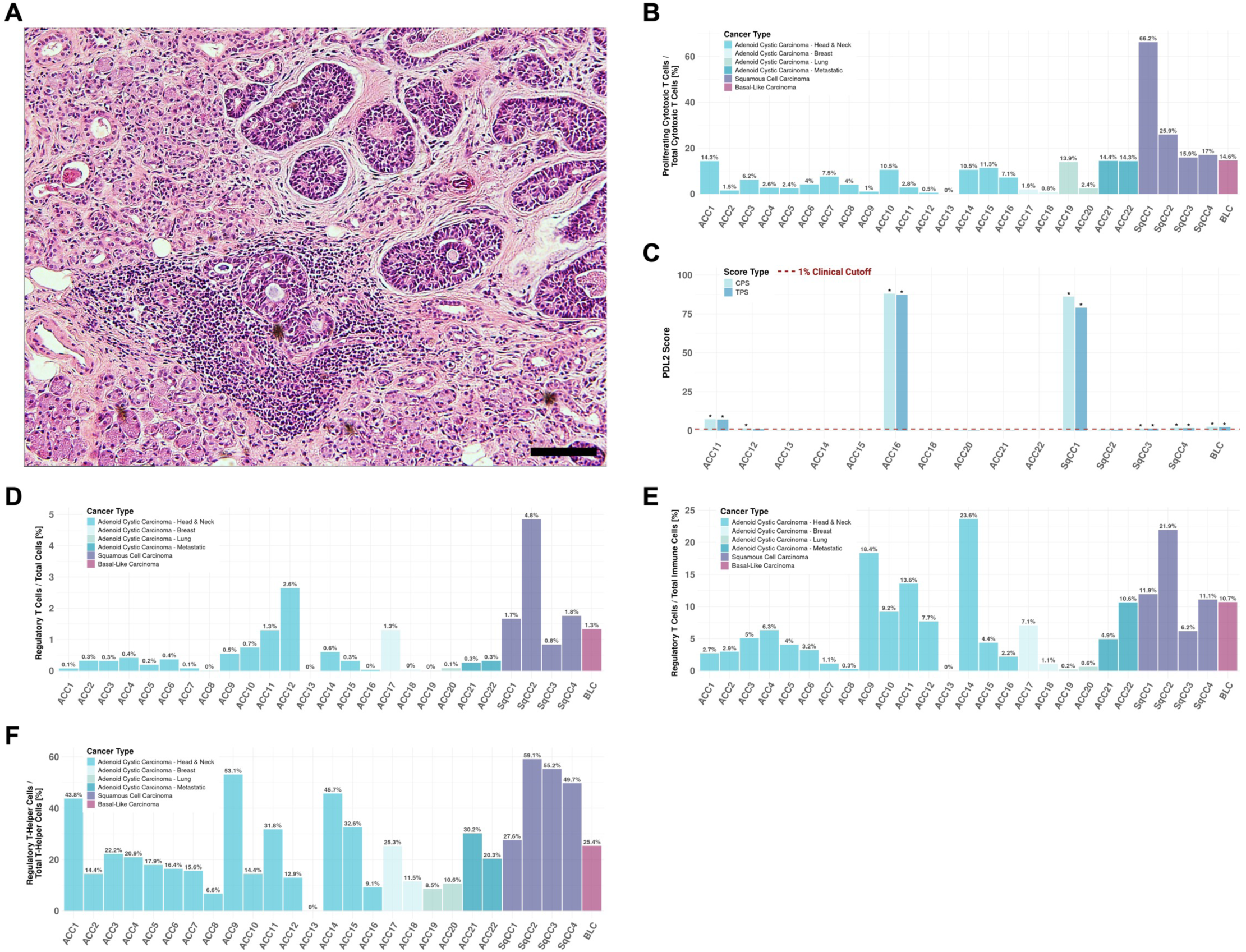
Further Immune Landscape Analysis. **A** Representative image of ACC and the adjacent normal salivary gland with tissue-resident immune cells. Image shown at 10x magnification. Scale bar, 100μm. **B** Percentage of proliferating cytotoxic T cells to total cytotoxic T cells. **C** Combined Positive Score (CPS) and Tumor Proportion Scores (TPS) for PD-L2 expression. Cases with scores of TPS >1% and CPS >1 are labeled with *. **D** Percentage of regulatory T cells to total cells in tumor area. **E** Percentage of regulatory T cells to total immune cells. **F** Percentage of regulatory T cells to total T-helper cells.

**Supplemental Figure 2.**
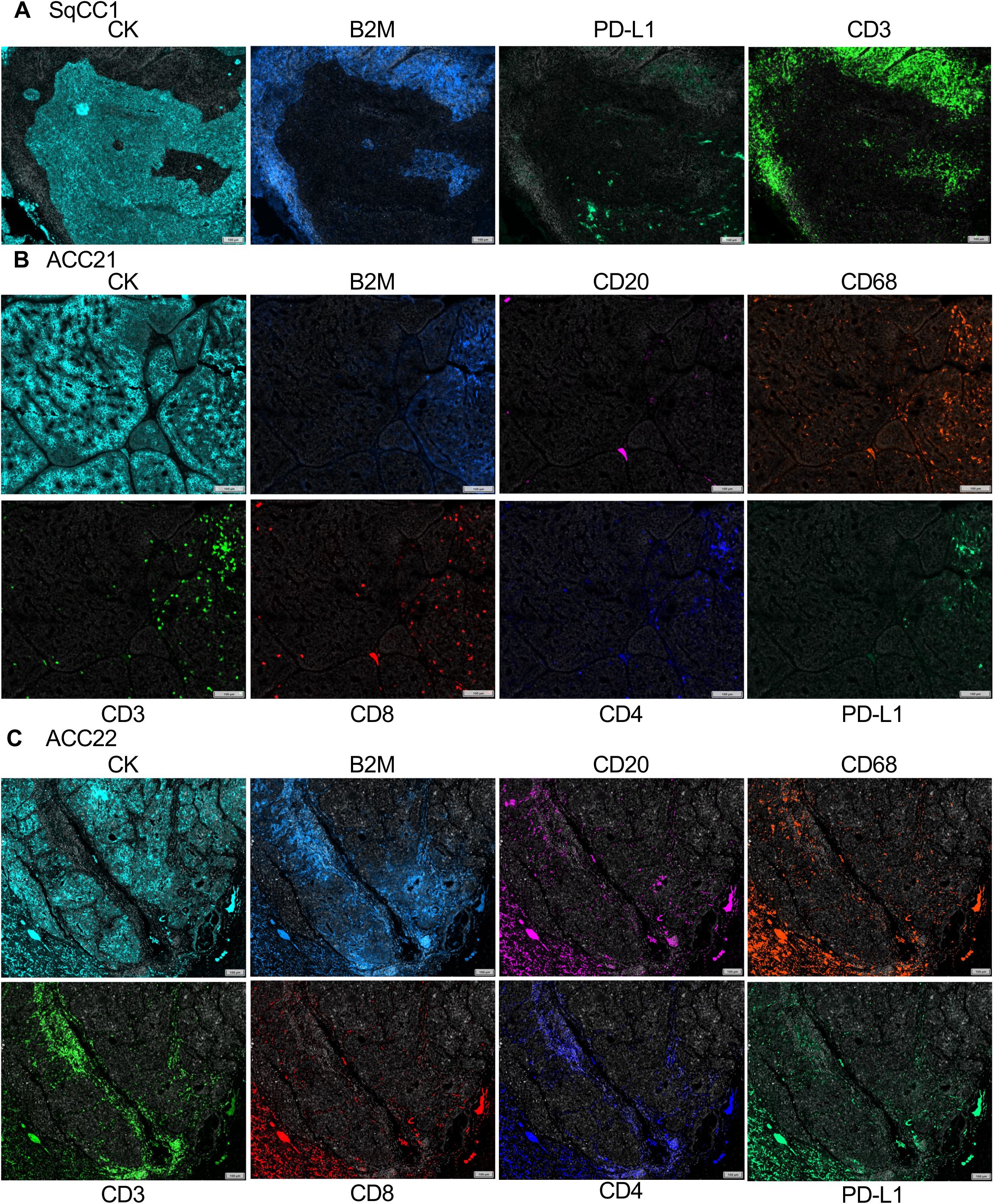
B2M Expression in Exceptional Cases. **A** Expression for CK, B2M, PD-L1, and CD3 in SqCC1, an HLA/B2M negative case in our cohort. **B, C** Expression for CK, B2M, CD20, CD68, CD3, CD8, CD4, and PD-L1 in ACC21 and ACC22, two focally B2M-positive, metastatic ACC cases. Scale bar, 100μm.

**Supplemental Figure 3.**
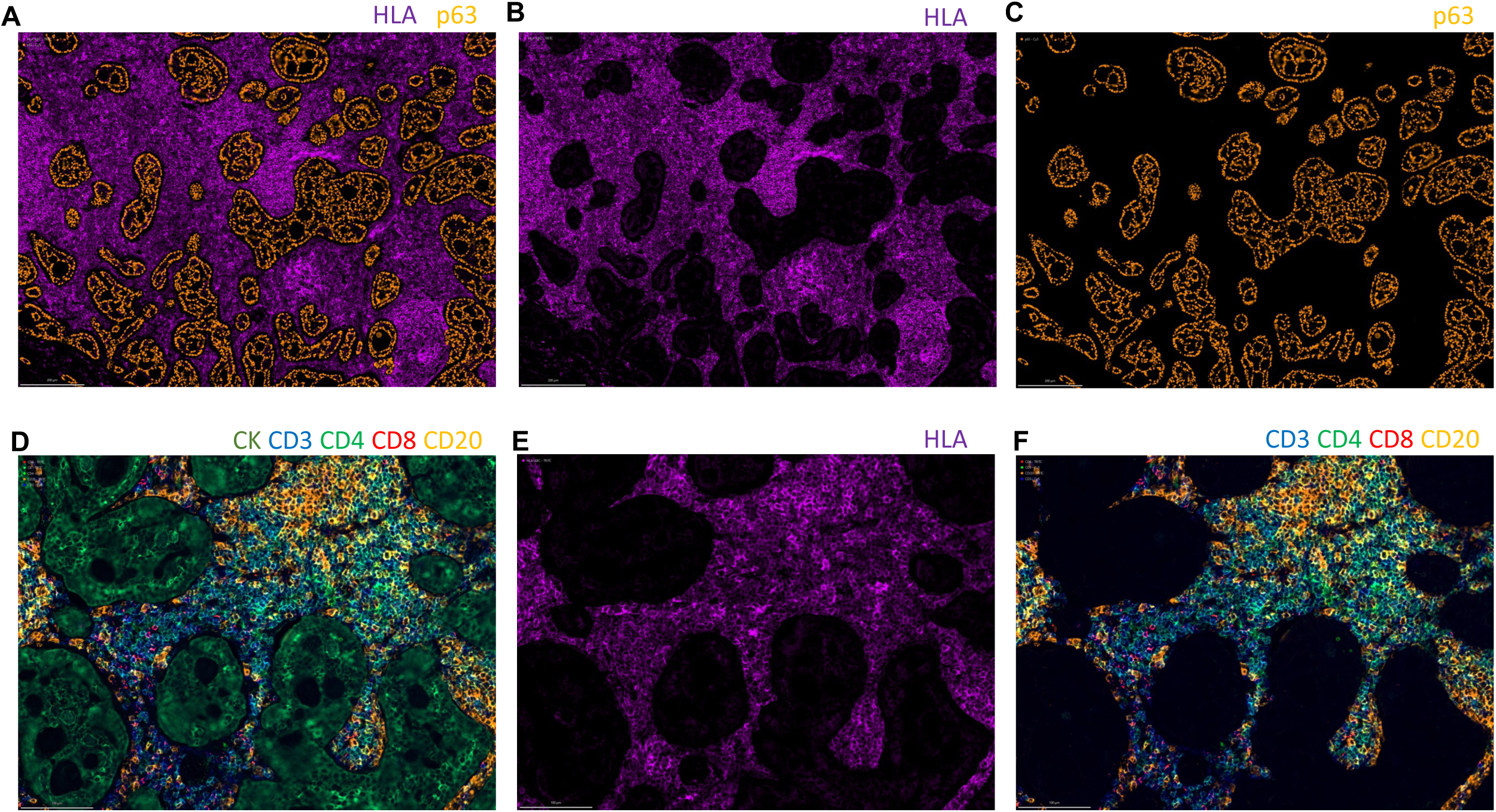
Representative images of immune landscape of an ACC case. **A** Overview images with Immunofluorescence staining of HLA and p63, and single channel images showing **B** HLA staining and **C** p63 staining. **D** Multiplex immunofluorescence image of CK, CD3, CD4, CD8 and CD20 staining, and image of the same area showing **E** HLA staining and **F** the immune cell markers CD3, CD4, CD8 and CD20 alone.

**Supplemental Figure 4.**
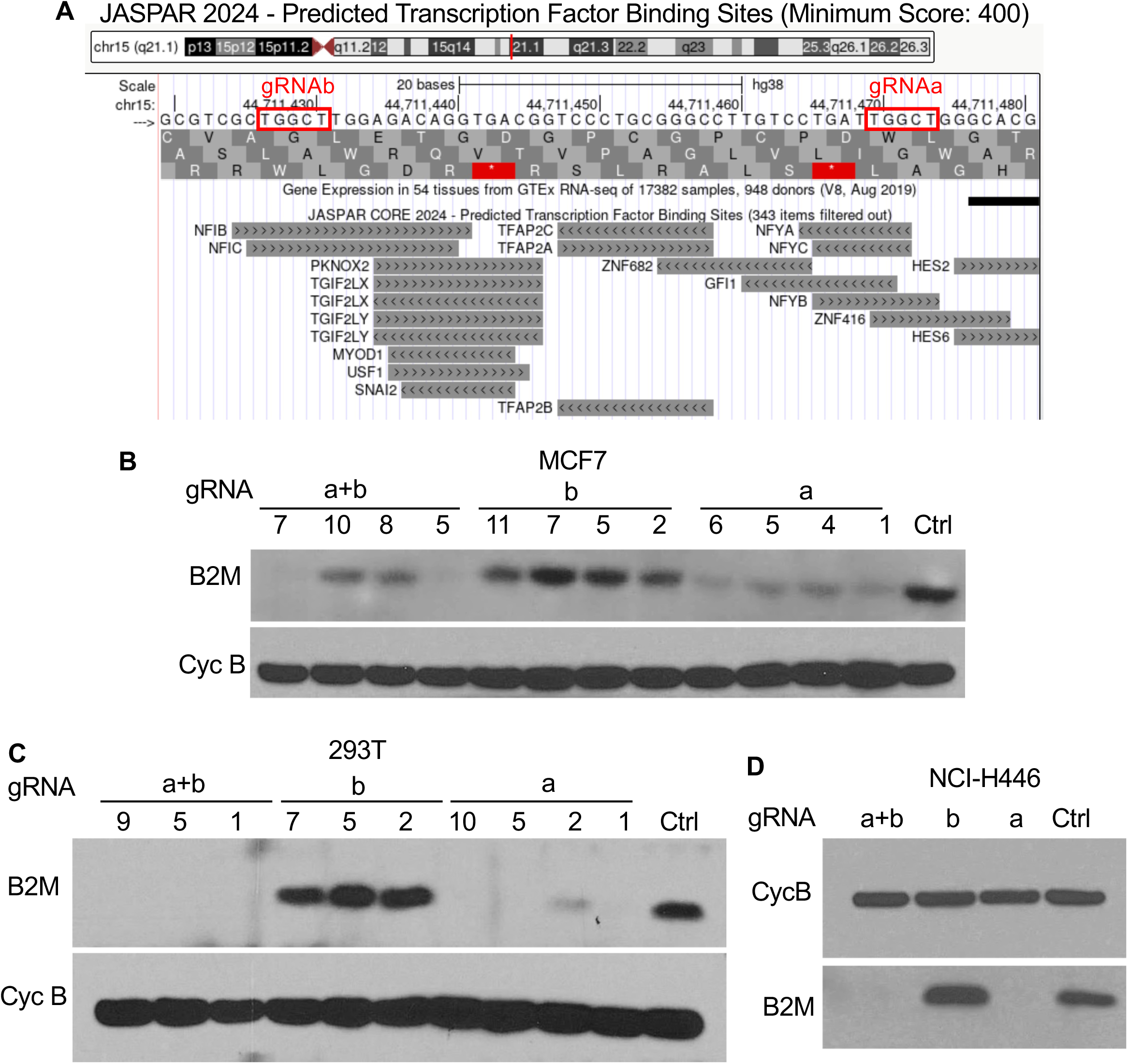
CRISPR-Cas9 Mediated Alteration of NFIB Binding Site at B2M Promoter. **A** Promoter region of B2M gene, showing the JASPAR 2024 predicted transcription factor binding sites with a minimum score of 400. The two target sequences of the gRNAa and gRNAb are highlighted. **B** Western blot showing expression of B2M in MCF7 cells stably expressing a control plasmid and single cell clones of cells expressing the gRNAa, gRNAb, and gRNAa+b plasmids. Cyc B acts as loading control. **C** Western blot showing expression of B2M in 293T cells stably expressing a control plasmid and single cell clones of cells expressing the gRNAa, gRNAb and gRNAa+b plasmids. Cyc B acts as loading control. **D** Western blot showing expression of B2M in NCI-H446 cells stably expressing a control plasmid and single cell clones of cells expressing the gRNAa, gRNAb and gRNAa+b plasmids. Cyc B acts as loading control. Cyc B, cyclophilin B.

**Supplemental Figure 5.**
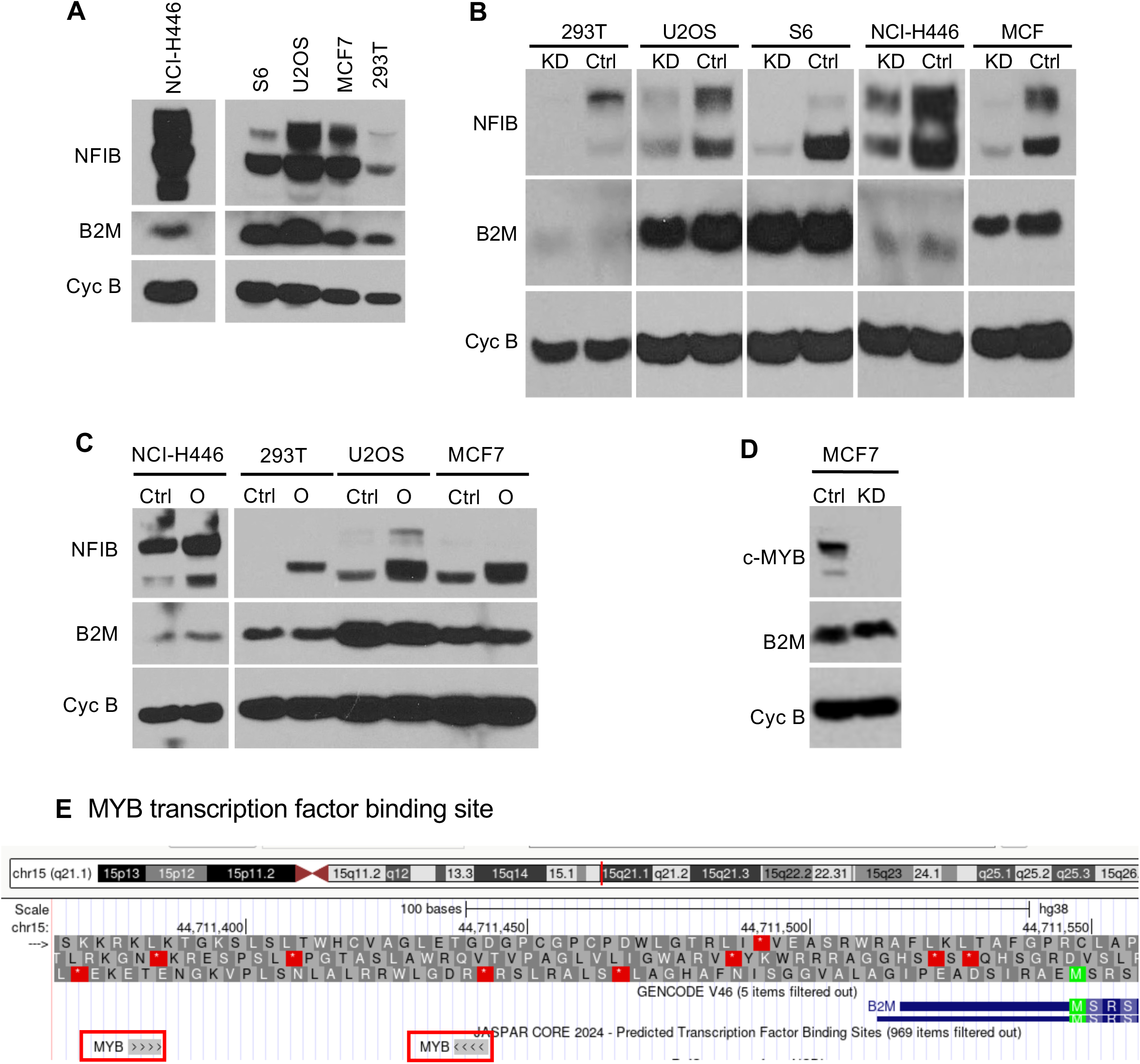
Effect of NFIB and MYB knockdown on B2M expression. **A** Western blot showing NFIB and B2M expression in wildtype cell lines NCI-H446, S6, U2OS, MCF7, and 293T. Cyc B acts as loading control. **B** Western blot showing NFIB and B2M expression in the cell lines NCI-H446, S6, U2OS, MCF7, and 293T transduced with a control shRNA or a NFIB knockdown shRNA. Cyc B acts as loading control. **C** Western blot showing NFIB and B2M expression in the cell lines NCI-H446, S6, U2OS, MCF7, and 293T transduced with a control plasmid or a NFIB overexpression plasmid. Cyc B acts as loading control. **D** Western blot showing c-MYB and B2M expression in MCF7 cells transfected with a control siRNA or MYB siRNA. Cyc B acts as loading control. **E** Promoter region of B2M gene, showing the JASPAR 2024 predicted transcription factor binding sites of MYB (JASPAR Score 243 and 249). Ctrl, control; KD, knockdown; O, overexpression; Cyc B, cyclophilin B.

**Supplemental Figure 6.**
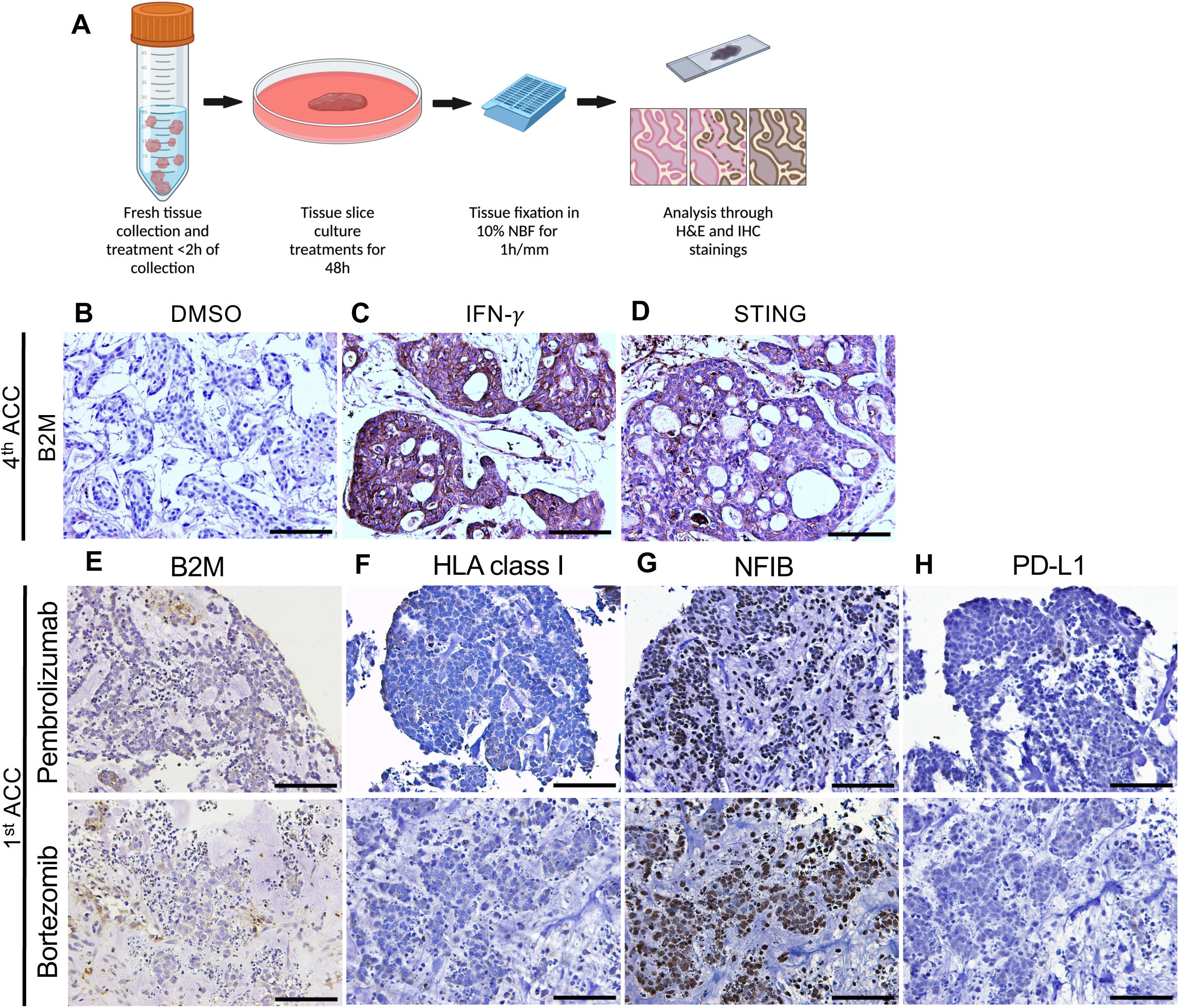
Treatment of Tissue Slice Cultures. **A** Workflow showing the collection and processing of fresh ACC tissue for 48h short-term cultures. Tissue is fixed for 1h/mm in 10% neutral buffered formalin (NBF) before embedding and processing for H&E, IHC, and IF analysis. **B-D** IHC staining for B2M and HLA class I of the 4^th^ ACC slice culture treated with DMSO (control), IFN-! or STING agonist. **E-H** IHC staining for B2M, HLA class I, NFIB, and PD-L1 of ACC tissues treated with pembrolizumab or bortezomib. Representative images shown at 20x magnification. Scale bar, 100μm.

**Supplemental Table 1.** Fusion Status of ACC Cohort. RNA-based fusion detection on 18 of the 21 ACC patients in our cohort found 13 fusion-positive cases. Ten cases showed a MYB-NFIB fusion, while the other 3 showed a MYBL1-NFIB fusion.

| sample | fusion | breakpoint1 | breakpoint2 | split_reads1 | split_reads2 | discordant_mates | coverage1 | coverage2 | confidence | reading_frame |
| --- | --- | --- | --- | --- | --- | --- | --- | --- | --- | --- |
| ACC1 | MYB:NFIB | chr6:135203324 | chr9:14084729 | 21 | 5 | 0 | 1163 | 10947 | high | out-of-frame |
| ACC4 | MYB:NFIB | chr6:135196002 | chr9:14102510 | 91 | 76 | 4 | 18790 | 15378 | high | in-frame |
| ACC5 | MYB:NFIB | chr6:135199050 | chr9:14088326 | 2 | 32 | 0 | 2896 | 8418 | high | in-frame |
| ACC7 | MYB:NFIB | chr6:135196002 | chr9:14102510 | 5 | 3 | 0 | 6192 | 13762 | high | in-frame |
| ACC11 | MYB:NFIB | chr6:135194460 | chr9:14088326 | 55 | 148 | 24 | 17450 | 22200 | high | in-frame |
| ACC12 | MYB:NFIB | chr6:135194460 | chr9:14088326 | 6 | 33 | 0 | 13974 | 34765 | high | in-frame |
| ACC13 | MYBL1:NFIB | chr8:66566684 | chr9:14116346 | 12 | 108 | 2 | 1366 | 4141 | high | in-frame |
| ACC14 | MYBL1:NFIB | chr8:66592440 | chr9:14102510 | 10 | 24 | 0 | 5303 | 4603 | high | in-frame |
| ACC17 | MYB:NFIB | chr6:135203324 | chr9:14102510 | 93 | 37 | 0 | 10442 | 14296 | high | in-frame |
| ACC20 | MYB:NFIB | chr6:135196002 | chr9:14088326 | 40 | 96 | 1 | 9944 | 20456 | high | in-frame |

| sample | fusion | RNA-based variants |
| --- | --- | --- |
| ACC2 | MYBL1 → NFIB | - |
| ACC8 | - | ALK p.Gly880Glu; FGFR2 p.Lys164Met |
| ACC15 | LOC100130691 → NFE2L2; MYB → INTERGENIC → PID1 | - |
| ACC19 | MYB p.Val314Leu | MYB → NFIB |
| ACC21 | - | - |
| ACC22 | - | - |

**Supplemental Table 2.** List of Primary Antibodies Used on the COMET mIF Platform. Antibody dilutions, supplier, catalogue number, and clone ID indicated for each antibody used in the panel.

| Antibody | Dilution | Supplier | Catalogue No | Clone ID |
| --- | --- | --- | --- | --- |
| **Beta-2-microglobulin** | 1:500 | Sigma-Aldrich, St. Louis, Missouri | HPA006361 | Polyclonal |
| **CD20** | 1:100 | Cell Marque, Rocklin, CA | 120M-86 | L26 |
| **CD3** | 1:500 | Cell Marque, Rocklin, CA | 103R-96 | MRQ-39 |
| **CD68** | 1:50 | Thermo-Fisher Scientific, Waltham, MA | MA5-13324 | KP1 |
| **CD4** | 1:400 | Abcam, Cambridge, UK | ab133616 | EPR6855 |
| **CD8** | 1:50 | Biorad, Hercules, CA | MCA1817 | 4B11 |
| **c-Myb** | 1:100 | Abcam, Cambridge, UK | ab45150 | EP769Y |
| **Cytokeratin 5** | 1:2500 | Thermo-Fisher Scientific, Waltham, MA | MA5-15347 | 3E251 |
| **FoxP3** | 1:50 | Thermo-Fisher Scientific, Waltham, MA | MA5-16365 | SP97 |
| **HLA1-ABC** | 1:1000 | Abcam, Cambridge, UK | ab70328 | EMR8-5 |
| **Ki67** | 1:25 | Agilent Technologies, Lexington, MA | M724001-2 | MIB-1 |
| **NFIB** | 1:100 | Abcam, Cambridge, UK | ab186738 | EPR14122 |
| **P63** | 1:2500 | Abcam, Cambridge, UK | ab124762 | EPR5701 |
| **Pan-cytokeratin** | 1:50 | Agilent Technologies, Lexington, MA | M351501-2 | AE1/AE3 |
| **PD-1** | 1:500 | Abcam, Cambridge, UK | ab137132 | EPR4877(2) |
| **PD-L1** | 1:100 | GenomeMe, Richmond, BC, Canada | IHC411-1 | IHC411 |
| **PD-L2** | 1:100 | Biotechne, Minneapolis, MN, USA | MAB-1224 | 176611 |
| **Smooth muscle actin** | 1:10000 | Abcam, Cambridge, UK | ab7817 | 1A4 |
| **SOX2** | 1:400 | Thermo-Fisher Scientific, Waltham, MA | MA1-014 | 20G5 |

**Supplemental Table 3.** Marker Combination for Cell Classification. The left column indicates the cell type or the cell subtype, while the right column shows the markers that need be above the threshold for the cells to be classified.

| Cell type | Positive markers |
| --- | --- |
| Epithelial | CK+ or CK+/PD-1+ |
| Epithelial: Proliferating | CK+/Ki67+ or CK+/Ki67+/PD-1+ |
| T-Helper | CD3+/CD4+ |
| T-Helper: Regulatory | CD3+/CD4+/FoxP3+ |
| T-Helper: on Epithelial | CD3+/CD4+/CK+ |
| T-Helper: Regulatory/on Epithelial | CD3+/CD4+/FoxP3+/CK+ |
| Cytotoxic T | CD3+/CD8+ |
| Cytotoxic T: Proliferating | CD3+/CD8+/Ki67+ |
| Cytotoxic T: on Epithelial | CD3+/CD8+/CK+ |
| Cytotoxic T: Proliferating/on Epithelial | CD3+/CD8+/Ki67+/CK+ |
| Cytotoxic T: Exhausted | CD3+/CD8+/PD-1+ |
| Cytotoxic T: Exhausted/on Epithelial | CD3+/CD8+/PD-1+/CK+ |
| Macrophage | CD68+ or CD4+/CD68+ |
| Macrophage: Proliferating | CD68+/Ki67+ or CD4+/CD68+/Ki67+ |
| Macrophage: on Epithelial | CD68+/CK+ or CD4+/CD68+/CK+ |
| Macrophage: Proliferating/on Epithelial | CD68+/CK+/Ki67+ or CD4+/CD68+/CK+/Ki67+ |
| Macrophage: Suppressed | CD68+/PD-1+ or CD4+/CD68+/PD-1+ |
| Macrophage: Suppressed/on Epithelial | CD68+/CK+/PD-1+ or CD4+/CD68+/CK+/PD-1+ |
| B Cell | CD20+ |
| B Cell: Proliferating | CD20+/Ki67+ |
| B Cell: on Epithelial | CD20+/CK+ |
| B Cell: Proliferating/on Epithelial | CD20+/CK+/Ki67+ |

